# Mapping genome-wide RNA-RNA and RNA-DNA interactions in nuclear hubs of male and female *Drosophila* cells

**DOI:** 10.64898/2026.08.02.742296

**Authors:** Sarah Gunasekera, Megan Carlson, Mukulika Ray, Erica Larschan

**Affiliations:** Computational Biology, Brown University, Providence, RI, USA; Molecular Biology, Cellular Biology, and Biochemistry Department, Brown University, Providence, RI, USA; Biology Department, University of Massachusetts Boston, Boston, Massachusetts, USA

## Abstract

Nuclear hubs regulate diverse nuclear processes by recruiting specific factors to the correct genomic loci at the correct time, ensuring the fidelity of transcription and RNA processing. However, how nuclear hubs are targeted to the genomic regions where they function remains poorly understood. One major challenge is capturing the higher-order RNA and DNA contacts that occur within these hubs. Here, we use RNA-DNA Split-Pool Recognition of Interactions by Tag Extension (RD-SPRITE) to map RNA-RNA and RNA-DNA contacts within nuclear hubs for the first time in Drosophila male (S2) and female (Kc167) cells. We observed higher-order hub structures in both cell types and identified several long non-coding RNAs (lncRNAs) that associate with spliceosomal RNAs. Some of these associations are sex-biased, including those of lncRNAs *CR34335* and *CR40469*, which have known roles in development and regeneration. We also investigated how the essential transcription factor CLAMP organizes spliceosomal RNA and DNA contacts, because we previously showed that CLAMP regulates splicing and prevents the formation of cryptic spliced isoforms. Our results suggest that CLAMP limits aberrant contacts between snRNAs and chromatin genome-wide, consistent with its role in preventing cryptic splicing. Our RD-SPRITE data also provide a valuable resource for studying other nuclear processes, such as miRNA-mediated silencing, nucleolar functions of snoRNAs, and Cajal body functions of scaRNAs. To facilitate broad community use, we developed Fly-SPRITE, a user-friendly computational platform that allows researchers to explore top RNA interactors and RNA-RNA associations and to identify the DNA targets of specific RNAs of interest.

## Introduction

The nucleus is organized into molecular compartments that coordinate functionally related RNA, DNA, and protein interactions into dense three-dimensional hubs. Within each of these compartments, biomolecules assemble within self-condensed regions in the nucleus to coordinate specific regulatory processes (Pombo & Dillon, 2015). For example, nucleolar hubs contain protein and RNA factors that catalyze the formation of ribosomal RNA and regulate ribosome biogenesis (Pelletier et al. 2018; Schöfer & Weipoltshammer, 2018), and spliceosomal compartments, or nuclear speckles, contain local concentrations of splicing factors and nascent transcripts that participate in RNA splicing (Belmont, 2022; Lamond & Spector, 2003). The high concentration of regulatory factors within compartments increases the efficiency of gene regulation by coordinating the association of co-regulated genes and the simultaneous processing of RNA. Mapping the molecular interactions that drive these processes, and how they coordinate with each other is therefore crucial to understand how molecular interactions within hubs drive the specificity of gene regulation.

Studies of molecular hubs in *Drosophila melanogaster* have explored transcription factor activity, enhancer-promoter interactions, and higher-order chromatin structures such as topologically associating domains (TADs) (Furlong & Levine, 2018; Mir et al., 2018). Together, this work has shown how proteins and DNA interact in 3D space to control precise gene expression. But, fewer studies have examined how RNAs work with proteins and DNA to shape hub formation, structure, and activity. Notable exceptions include the roX1 and roX2 long non-coding RNAs (lncRNAs) in *Drosophila* dosage compensation (Conrad and Akhtar, 2012; Frank and Baker 1999) and the hsr-omega nuclear (hsrω) lncRNA in hnRNP-containing omega speckles (Prasanth et al. 2000; Singh and Lakhotia 2015). In both cases, the RNAs play structural and regulatory roles within its hub. Whether other RNAs act in similar ways in *Drosophila* hubs has not been tested systematically.

The development of proximity ligation methods such as ChAR-seq (Bell et al., 2018) has made it possible to map the localization of RNA species throughout the genome. These methods can identify where a specific RNA species is located and whether it belongs to a ribonucleoprotein (RNP) complex to deduce its regulatory function. However, proximity ligation methods are limited in their ability to capture the full combinatorial complexity of molecular hubs, as they cannot simultaneously resolve multiple RNA and DNA species and association within the same condensate. This limitation is especially challenging for large or dynamic complexes that have multiple conformations and consist of many molecular partners. The RNA & DNA SPRITE (RD-SPRITE) method (Quinodoz et al., 2021) overcame this limitation by capturing thousands of higher-order RNA and DNA contacts, revealing multi-way interactions within compartments including the co-clustering of small nuclear RNAs (snRNAs) within spliceosomal complexes (Bhat et al., 2024; Quinodoz et al., 2021). Because RNAs that share function tend to be localized together, these interaction patterns can help generate hypotheses about the roles of poorly characterized RNAs. For these reasons, RD-SPRITE is a powerful tool for studying the molecular organization and function of compartments throughout the cell. Despite this potential, its application has so far been limited to human and mouse samples, and comparisons across different model organisms and sexes have yet to be explored.

Here, we implemented the RD-SPRITE method for the first time in *Drosophila* male (S2) and female (Kc167) cells to define pairwise and multi-way RNA-RNA and RNA-DNA interactions within nuclear compartments. Using this approach, we identified RNA and chromatin compartments organized around spliceosomal, ribosomal, and other nuclear RNAs. Within spliceosomal multi-way interactions, we found several lncRNAs that interact abundantly with snRNAs. These lncRNAs have known functions in development (Camilleri-Robles et al., 2024) but have not yet been characterized for a role in *Drosophila* alternative splicing (AS). In human cells, lncRNAs such as MALAT1 regulates AS by modulating how splicing factors associate with precursor messenger RNAs (pre-mRNAs) (Tripathi et al., 2010). However, the mechanisms by which lncRNAs regulate AS in *Drosophila* are poorly understood. Beyond splicing, our male and female RD-SPRITE datasets provide a valuable resource for investigating other processes in *Drosophila*, such as nucleolar functions of snoRNAs and Cajal body functions of scaRNAs. To facilitate broad community use, we developed a computational platform, Fly-SPRITE, that enables *Drosophila* researchers to explore, for a specific RNA, its top interactors, sex-biased RNA-RNA interactions, and DNA targets in our male and female datasets. The platform also allows simultaneous analysis of multiple RNAs.

Previous studies have also identified the *Drosophila* DNA binding protein, CLAMP (Chromatin-linked Adapter for MSL Proteins) as a key regulator of alternative splicing that prevents cryptic splicing events (Ray et al. 2023; Ray et al. 2026). CLAMP associates with protein and RNA factors that regulate splicing (Kaye et al., 2018), binds intronic regions as frequently as promoters (Kaye et al., 2018), and its binding sites evolved from intronic polypyrimidine tracts (Quinn et al., 2016) – all hallmarks of splicing regulatory activity. Furthermore, CLAMP regulates sex-specific splicing (Ray et al., 2023), directly binds several snRNAs (Ray et al., 2026), and prevents cryptic splicing (Ray et al., 2026). Cryptic splicing occurs when the spliceosome recognizes and processes splice sites that are normally ignored. Given the established role for CLAMP in regulating sex-specific splicing, and RD-SPRITE’s ability to capture higher-order contacts among spliceosomal RNAs, we also performed RD-SPRITE in *Drosophila* male (S2) and female (Kc167) cells following depletion of CLAMP. We identified several snRNA species that showed a global increase in chromatin occupancy upon CLAMP depletion, consistent with the model in which CLAMP prevents cryptic splicing activity by blocking aberrant association of spliceosomal hubs with chromatin.

Together, we provide the first mapping of RNA-RNA and RNA-DNA contacts within nuclear hubs in *Drosophila* using the RD-SPRITE approach. We make our RD-SPRITE male and female *Drosophila* datasets publicly available for researchers to explore RNA targets of interest through our Fly-SPRITE platform.

## Results

We implemented RD-SPRITE in the well-established male (S2) and female (Kc167) *Drosophila* cell lines that are transcriptionally similar and derive from hemocyte lineage (Drewell et al. 2024; Kloranos et al. 2023) to identify multiple RNAs and DNAs that interact with each other simultaneously (Figure 1a). As defined by the RD-SPRITE method, we used a split-and-pool barcoding approach which tags interacting RNAs and DNAs with a unique shared barcode (Quinodoz et al., 2021). We then leveraged the existing RD-SPRITE pipeline (Quinodoz et al., 2021) and configured it for *Drosophila* genomes to process sequenced reads separately for male and female cells. We identified the interacting groups, or clusters, of RNAs and DNAs based on its unique barcode. Cluster sizes can vary between 1 and more than 1,000 reads of RNAs and DNAs. After combining technical replicates, we removed singletons and clusters with more than 1,000 reads. For Experiment A, we retained ∼920,000 and ∼500,000 RNA and DNA reads combined from the male 2.5% and 5% libraries and retained ∼600,000 and ∼400,000 reads combined from the female 2.5% and 5% libraries (Table S1). We found that most clusters contained in male and female RD-SPRITE cluster files contained 2-10 reads (75-86%) (Figure S1). All analyses in Figure 1 and Figure 2 used interaction data from Experiment A, 2.5% male and female libraries.

**Figure 1:**
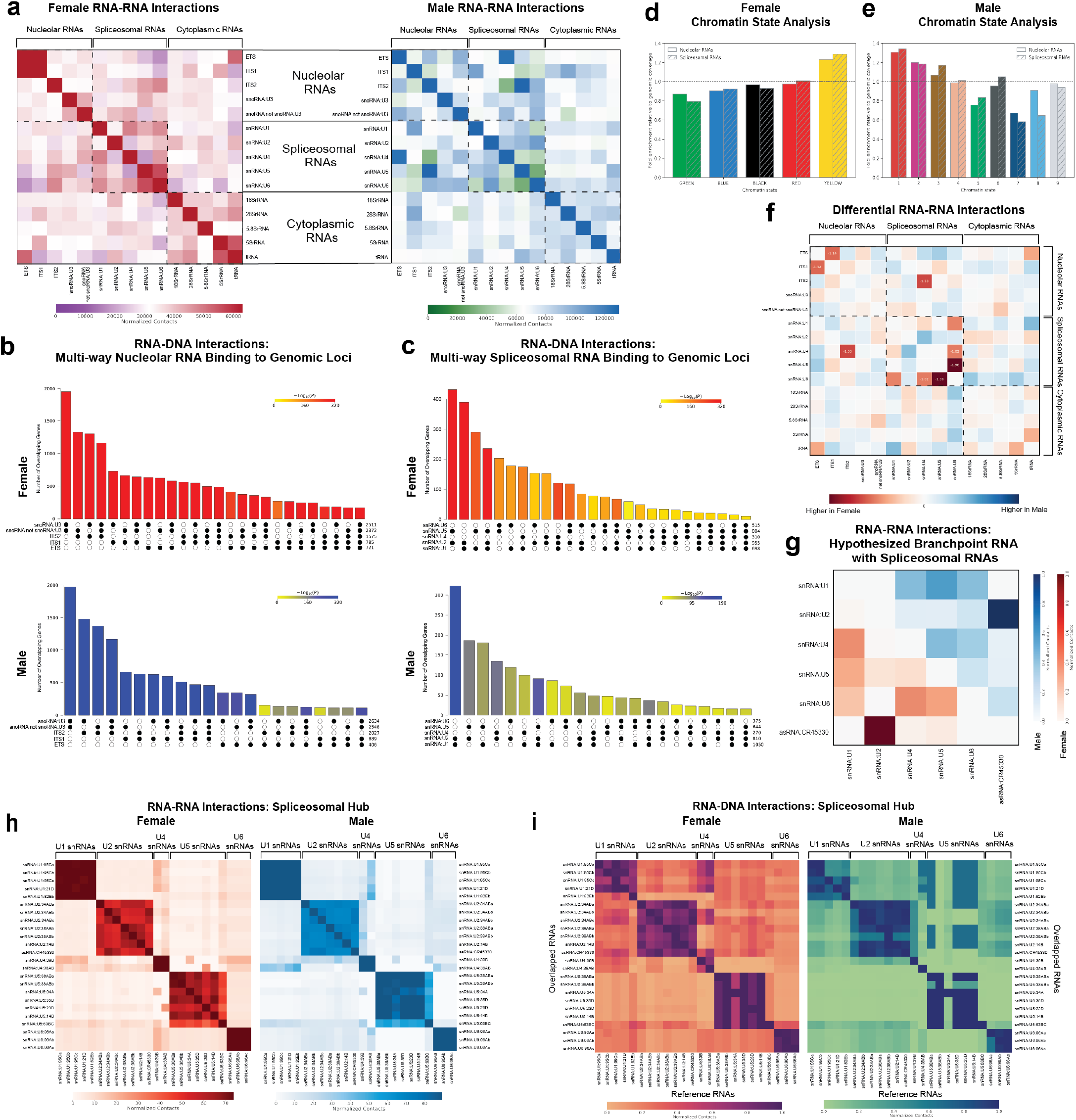
Higher-order RNA and DNA contacts among functionally related RNAs in *Drosophila melanogaster* revealed by RD-SPRITE. All analyses in Figure 1 used the 2.5% male (S2) and female (Kc167) control libraries from Experiment A. **(a)** Heatmap showing pairwise RNA-RNA interactions in female (left) and male (right) cells. Boxes denote functional groups of plotted nucleolar, spliceosomal, and cytoplasmic RNAs. Only the major spliceosomal RNAs are plotted within the spliceosomal group. **(b-c)** Overlap of genes bound with nucleolar RNA (b) and major spliceosomal RNA (c) in female (top, red) and male (bottom, blue) cells. The matrix of solid and empty circles at the bottom depicts the “presence” or “absence” of that RNA’s gene set in each intersection. The numbers to the right of the matrix are gene-binding set sizes for each RNA. Intersections were assembled using the SuperExactTest (Wang et al. 2015) and are ordered by size. Only intersections of at least 100 genes (b) or 10 genes (c) are shown. Darker red (female) or blue (male) bars indicate smaller P values. **(d-e)** Chromatin occupancy of RNAs involved in nucleolar and spliceosomal function in each chromatin state, shown as fold change enrichment of that state’s genome-wide percentage in female (d) and male (e) cells. **(f)** Heatmap showing pairwise differential contact RNAs involved in nucleolar, spliceosomal, and cytoplasmic function calculated from male and female RNA-RNA interaction profiles. Normalized log2 fold change in RNA-RNA interaction frequency between male and female cells. Red (left) indicates interactions higher in female cells, and blue (right) indicates interaction higher in male cells. Values are shown for interactions with |log2 fold change| > 2. **(g)** Heatmap showing pairwise RNA-RNA interactions between major spliceosomal RNAs and antisense lncRNA *CR45330* in female (bottom; red) and male (top; blue) cells. For females and males, the red/white and blue/white color bar, respectively, represent high and low normalized contacts. **(h)** Heatmap showing pairwise RNA-RNA interactions between major spliceosomal RNAs in female (left) and male (right) cells. For females and males, the red/white and blue/white color bar, respectively, represents high and low snRNA RNA-RNA normalized contacts. **(i)** Heatmap showing overlapped RNA-DNA binding sites between major spliceosomal RNAs in female (left) and male (right) cells. For females and males, the purple/orange and blue/green color bars, respectively, represent high and low correlations of pairwise RNA localization to DNA.

**Figure 2:**
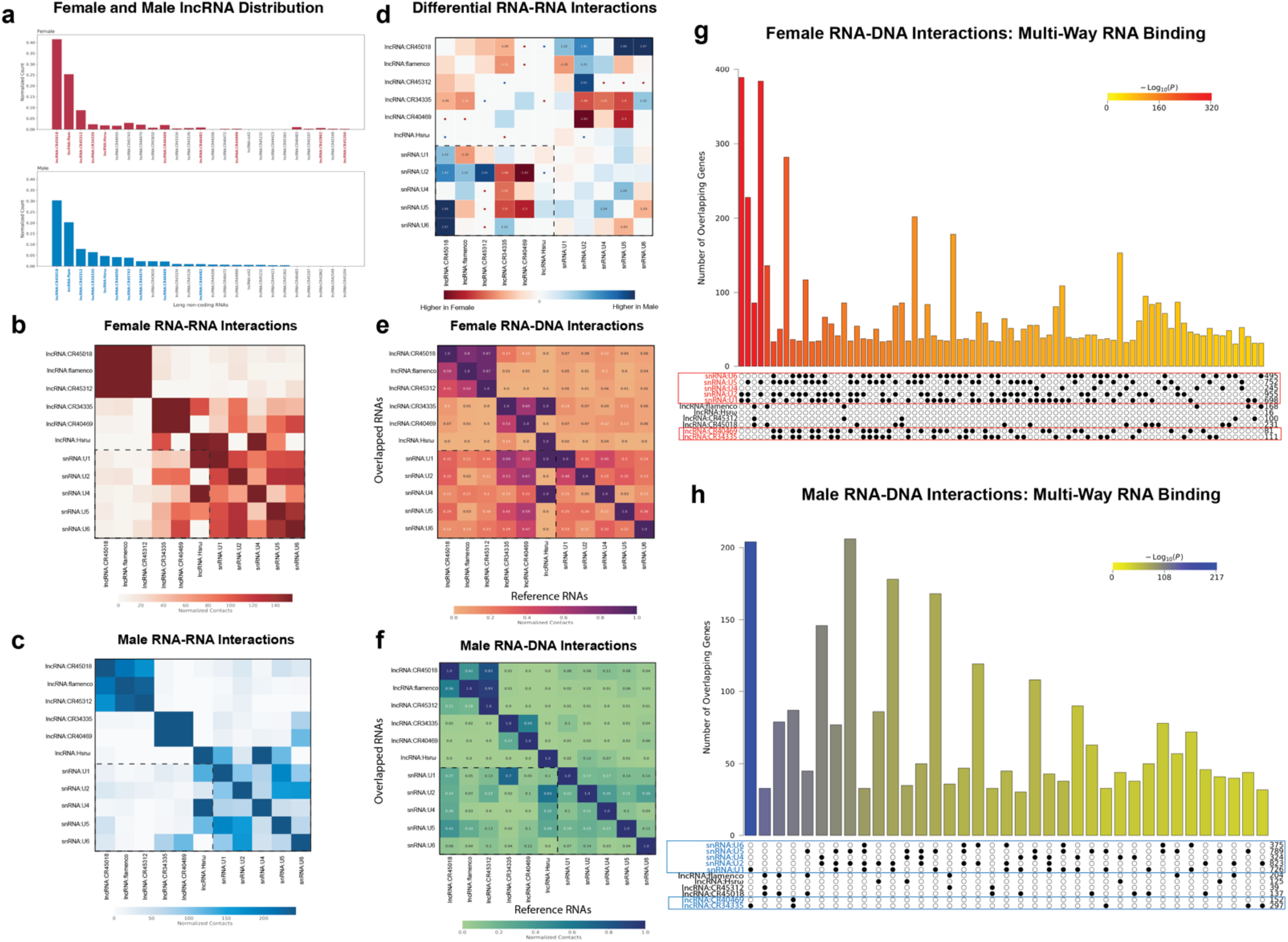
Long non-coding RNAs display enriched RNA and DNA contacts with major spliceosomal RNAs. All analyses in Figure 2 used the 2.5% male (S2) and female (Kc167) control libraries from Experiment A. **(a)** Top 20 most abundant lncRNAs within female (red) and male (blue) RD-SPRITE clusters. The top 10 lncRNAs that share the most snRNA interaction targets are bolded. **(b-c)** Heatmap showing pairwise RNA-RNA interactions between major spliceosomal RNAs and the top 6 snRNA interacting lncRNAs (dotted box) in female (b) and male (c) cells. **(d)** Heatmap showing pairwise differential contacts calculated from male and female RNA-RNA interaction profiles. Normalized log2 fold change in RNA-RNA interaction frequency between male and female cells. Red (left) indicates interactions higher in female cells, and blue (right) indicates interactions higher in male cells. Values are shown for interactions with |log2 fold change| > 2. Dots mark interactions detected in only one sex, colored by the sex in which they were detected. **(e-f)** Heatmap showing pairwise RNA-RNA interactions at shared DNA genomic regions between major spliceosomal RNAs and 6 of the top 10 snRNA-interacting lncRNAs (dotted box) in female cells (e). The same 6 lncRNAs are plotted in male cells for comparison (f). **(g-h)** Overlap of genes bound with abundant lncRNAs (*flamenco*, *Hsrω, CR45312, CR45018, CR40469, CR34335*) and major spliceosomal RNA in female (g) and male (h) cells. The matrix of solid and empty circles at the bottom depicts the “presence” or “absence” of that RNA’s gene set in each intersection. The numbers to the right of the matrix are gene-binding set sizes for each RNA. Darker red (g) or blue (h) bars indicate smaller P values. Intersections were assembled using the SuperExactTest (Wang et al. 2015) and are ordered by P value (smaller to larger). Only intersections of at least 30 genes are shown.

### 1.1 A subset of ncRNAs with shared biological function are present within the same spatial hubs within the nucleus

To characterize pairwise interactions among multiple RNAs, we defined RNA-RNA and RNA-DNA interactions from our male and female RD-SPRITE clusters. The RNAs selected for this analysis are non-coding and part of three broad groups each with distinct functions: the nucleolar group, spliceosomal group, and cytoplasmic group. The nucleolar groups consist of RNAs involved in ribosomal RNA (rRNA) processing, the spliceosomal group regulates RNA splicing, and the cytoplasmic group regulates translation. We focused on normalized pairwise contact frequencies for RNA-RNA interactions and normalized RNA-DNA contact frequencies derived from overlapping BED file entries. We found that RNAs that share the same broad biological functions share RNA interactions (Figure 1a) and bind to shared genomic loci (Figure S2a). However, several RNAs also interact with RNAs from other functional groups and share genomic contacts with them. For example, the U3 snoRNA, a nucleolar RNA that is required for ribosome biogenesis and canonically pairs with pre-rRNA (Cass et al. 1990), shares both RNA and DNA interactions with snRNAs (Figure 1a; S2a). This differs from the RNA and DNA interaction profiles generated from mammalian RD-SPRITE datasets, which showed distinctly higher pairwise interaction frequencies among RNAs within functional groups but not between groups (Quinodoz et al., 2021).

We then asked whether RNAs within the same functional group also participate in multi-way interactions to form higher-order structures. To measure statistical significance of the multi-way RNA interactions among ribosomal, spliceosomal, and cytoplasmic RNAs, we implemented the multi-way contact score (Quinodoz et al., 2021) in male and female cells. This metric is calculated through comparing the frequency of contacts between three or more RNAs to the expected frequency if these RNAs were randomly distributed. We observed a highly significant number of multi-way RNA interactions within the nucleolar, cytoplasmic, and spliceosomal RNA groups across a range of higher-order structure sizes (i.e. k-mers) (Table S2). We also measured multi-way RNA interactions on chromatin using the SuperExactTest (Wang et al. 2015). Using this method, we overlapped gene lists bound by each RNA and calculated the exact probability of multi-set intersections. We observed that at multiple higher-order structure sizes, the nucleolar and spliceosomal RNAs share multi-way interactions on chromatin within their functional group (Figure 1b; 1c). Consistent with prior approaches (Quinodoz, 2021), we define groups of RNAs exhibiting multi-way RNA interactions as “hubs.”

### 1.2 ncRNAs involved in rRNA processing share multi-way RNA and DNA interactions

The nucleolar hub is the site of ribosomal biogenesis which begins with the transcription of the ribosomal genes, and maturation of the 47S pre-ribosomal RNA (pre-RNA) in the nucleolus (Boisvert et al., 2007). The pre-rRNA matures through the cleavages of external and internal transcribed spacer regions (ETS and ITS) which is carried out by complexes containing small nucleolar RNAs (snoRNAs) together with associated proteins called small nucleolar ribonucleoproteins (snoRNPs) (Matera et al., 2007). SnoRNAs were initially discovered for their roles in the processing of rRNA, but they are also responsible for post-transcriptional modifications of other RNA species, including small nuclear RNAs (snRNAs), transfer RNAs (tRNA), and messenger RNAs (mRNAs) (Kass et al. 1990; Jady, 2003; Vitali & Kiss, 2019). SnoRNAs perform a variety of roles; therefore, we split the snoRNAs into two categories: (1) U3 snoRNA which specifically promotes pre-rRNA cleavage (Cass et al. 1990), and (2) all other snoRNAs that guide other functions and modifications (Huang et al. 2022). We found that nucleolar RNAs form multi-way RNA contacts with each other (Table S1) and exhibit shared colocalization to chromatin in male and female cells (Figure 1b). In both males and females, U3 snoRNA, other snoRNAs, and the ITS2 region colocalize with chromatin at the greatest number of genes, followed by the ITS1 region. The ETS region binds to the most shared genomic loci with U3 snoRNA and the other snoRNAs (Figure 1b). Overall, nucleolar RNAs exhibit multi-way interactions on the RNA-RNA and RNA-DNA interaction level, supporting evidence of nucleolar hub formation in male and female *Drosophila* cells.

### 1.3 ncRNAs involved in pre-mRNA splicing share multi-way RNA and DNA interactions

The spliceosomal hub is the site of RNA splicing mediated by the spliceosome ribonucleoprotein (RNP) complex. The spliceosome consists of major and minor spliceosomal complexes, responsible for splicing two types of introns: U2- and U12-type introns, respectively. The major spliceosomal unit, the U2 spliceosome, is formed through interactions between U1, U2, U4, U5, and U6 RNPs and RNAs (Burge et al., 1998), while the minor spliceosomal unit, U12 spliceosome, is formed through interactions between U11, U12, U4atac, U6atac, and U5 RNPs and RNAs (Hall and Padgett, 1996; Tarn & Steitz, 1996). The major spliceosome catalyzes the removal of more than 99% of all introns, whereas the minor spliceosome splices less than 1% of introns (Alioto, 2007). While RNAs belonging to the minor spliceosomal unit were identified in our male and female RD-SPRITE datasets, we omitted them from this analysis due to low read counts.

We found that the major spliceosome snRNAs form multi-way RNA contacts with each other in male and female cells (Table S1), and combinations of these RNAs colocalize with shared genomic regions (Figure 1c). In both males and females, U1, U2, and U5 snRNAs show the greatest degree of co-occupancy with chromatin (Figure 1c), sharing the largest number of genomic loci. Interestingly, colocalization of all five major spliceosomal snRNAs – U1, U2, U4, U5, and U6 – occurs at a lower frequency than pairwise or partial combinations (Figure 1c), suggesting that partial co-occupancy of individual snRNAs may be more common than complete spliceosome assembly at many genomic loci. Overall, these results suggest snRNAs exhibit multi-way interactions on the RNA-RNA and RNA-DNA interaction level, revealing preferences for specific combinatorial interactions among subsets of snRNAs within spliceosomal hubs in males and females.

### 1.4 Nucleolar and spliceosomal RNAs associate with transcriptionally active euchromatin regions of the genome in both sexes

To investigate the chromatin environments that nucleolar and spliceosomal RNAs contact, we compared their RNA-DNA contacts to the *Drosophila* S2 9-state (Kharchenko et al., 2011; Figure S3c) and the Kc167 5-state (Filion et al., 2010; Figure S3a) HMM chromatin model. These models bin the genome into distinct states by integrating dozens of modENCODE chromatin profiling datasets. Compared to the genome-wide distribution of chromatin states, nucleolar and spliceosomal RNA contacts were enriched in transcriptionally active chromatin (Figure 1d, e). In male cells, enrichment was highest in active states: state 1 (∼1.30 fold for nucleolar and spliceosomal RNAs), state 2 (∼1.20 fold) and state 3 (∼1.10 fold). In female cells, both groups were enriched in the active YELLOW state (∼1.20 fold). These active states occupy a small fraction of the genome: ∼9%, ∼13%, and ∼8% for active states 1, 2, and 3 in S2 cells (Figure S3d), and ∼19% for the active YELLOW state in Kc cells (Figure S3b). Together, these results suggest that spliceosomal and nucleolar RNAs contact transcriptionally active chromatin more than expected from the distribution of these states alone. A similar trend was observed for RNA contacts overall, suggesting that part of this enrichment may also reflect a general preference of RD-SPRITE for capturing contacts between RNA and accessible chromatin.

### 1.5 Sex-biased trends in nucleolar, spliceosomal, and cytoplasmic RNA interactions

Having identified higher-order RNA interactions within functional groups in both sexes as expected, we next investigated how these interactions may differ between males and females. Although male S2 and female KC167 cells have broadly similar transcriptional profiles (Drewell et al. 2024), previous studies have identified 2,588 differentially expressed genes between them (Kloranos et al. 2023) which may partly reflect sex-specific gene regulation. Sex-specific alternative splicing has also been shown in *Drosophila* (Ray et al., 2023). To reveal interaction level differences that may contribute to sex-biased expression, we calculated a differential matrix to compare differences between the male and female RNA-RNA interaction profiles among the nucleolar, spliceosomal, and cytoplasmic RNAs. As expected for nuclear hubs required for cell viability, only 4 RNA-RNA interaction pairs showed a sex-biased difference (Figure 1f; log2FC > 1). The only sex-biased interactions among the interaction pairs were female-enriched (Figure 1f). Due to the limited number of biological replicates that we could generate, we describe sex-biased interactions as trends that we observe in our data.

### 1.6 A non-snRNA species associates with spliceosomal U2 snRNAs

Among the RNA interactors with major spliceosomal RNAs, we identified an RNA (asRNA:CR45330) that interacts preferentially with the U2 snRNA compared to the other snRNAs in both males and females (Figure 1g). During the early stages of spliceosome assembly, U1 snRNP base pairs with the 5’ splice site while U2 snRNP recognizes and binds the intron branch point sequence, enabling the first catalytic step of pre-mRNA splicing (Ares and Weiser 1995). The antisense long non-coding RNA CR45330 (asRNA:CR45330) has been computationally predicted to be a component of the U2 snRNP on the basis of sequence similarity to the Rfam U2 snRNA family (RF00004), with predicted roles in branch point binding and branch site recognition (Kalvari et al., 2018; FlyBase, FBgn0266869). Consistent with this prediction, our results demonstrate that CR45330 asRNA exhibits enriched RNA-RNA interactions with U2 snRNA, suggesting that it may function in close association with the U2 snRNP. Therefore, we determined that a predicted component of the U2 spliceosome associates with it *in vivo*, opening a new area for future investigation.

### 1.7 The RNA and DNA interactions among spliceosomal snRNA variants reflect their organization into distinct snRNP complexes

The major spliceosomal snRNPs are each defined by their respective snRNA that is stably associated with a set of protein components that together drive pre-mRNA splicing. These snRNAs consist of multiple paralogous genes. Compared to vertebrates, *Drosophila melanogaster* has many fewer snRNA paralogs: five U1 genes, six U2 genes, three U4 genes, seven U5 genes, and three U6 genes (Lu and Matera, 2015). The other minor spliceosomal snRNAs are all expressed from single copy genes (Lu and Matera, 2015). All of the transcripts for each of the gene paralogs are found in our RD-SPRITE datasets; however, we omitted the minor snRNAs and the U4 snRNA variant, snRNA:U4:25F, due to low read depth.

When we examined the higher-order contacts of these snRNAs grouped by class – U1, U2, U4, U5, and U6 – we found that major snRNAs share RNA interactions (Figure 1a) and bind to shared genomic loci as expected (Figure 1c). To determine whether transcript variants from a specific snRNA class display distinct interaction structure, we generated RNA-RNA and RNA-DNA interaction profiles for all snRNA variants. In both sexes, transcript variants within the same snRNA class share RNA interactions (Figure 1h, S2b, S2c) and bind to shared genomic loci more with one another than with variants from other classes (Figure 1i), suggesting that class-specific function governs spatial organization. We also observed that certain snRNA transcripts exhibit different binding frequencies than their snRNA counterparts within the same class. Within the U5 snRNA family, 63BC U5 snRNA interacts with variants within the U1, U2, U4 snRNA families in both sexes (Figure 1h), consistent with prior work showing that 63BC U5 snRNA assembles into large, complex snRNPs that exclude other U5 paralogues (Chen et al., 2005). Our results extend this work by identifying the variants from other snRNA families that U5:63BC contacts. Furthermore, asRNA:CR45440 interacts predominantly with U2 snRNA variants (Figure 1h and 1i), which further reinforces its potential role in U2 snRNP complex function.

Having established that RD-SPRITE captures bona fide, class-organized snRNA substructure at both the RNA-RNA and RNA-DNA interaction levels, we next investigated how snRNA variant RNA-RNA interactions differ between sexes. We found that interactions among snRNAs within a class have no sex biased differences. However, interactions between the snRNA variants, snRNA:U4:39B, snRNA:U4:38AB, and snRNA:U5:23D, and U1 and U2 snRNAs exhibit sex-biased interactions (Figure S2d; log2FC > 1). Pairwise interactions that are observed in one batch but not the other are listed with a ‘dot’ but due to the limited number of biological replicates, these results are described as trends.

Together, these results indicate that higher-order spatial organization of functionally related ncRNAs occurs at the RNA interaction level and at shared genomic sites in *Drosophila*. Furthermore, within these broader hub interaction networks, distinct sub-structures emerge that may reflect more specialized roles within hub machinery.

### 2.1 Several long non-coding RNAs are present within RD-SPRITE clusters

Besides the small ncRNAs mentioned earlier, long non-coding RNAs (lncRNAs), a diverse group of ncRNAs more than 200 nucleotides in length, play important roles in the nucleus (Yu and Shan, 2016). LncRNAs associate with nuclear components, modify chromatin, and regulate gene expression at multiple levels, and many form nuclear hubs (Mattick et al., 2023; Hasenson et al., 2022; Takakuwa et al., 2023). We therefore examined lncRNA contacts, focusing on lncRNAs that contact RNAs with established roles in nuclear hubs that regulate gene expression. To identify the most abundant lncRNAs in male and female cells, we ranked all lncRNAs detected in our RD-SPRITE cluster files and selected the top 20 in each (Figure 2a). Among these lncRNAs, only a small subset have been functionally characterized: (1) *hsrω*, which forms nuclear speckles containing RNA-binding proteins involved in splicing that help regulate gene expression during heat stress (Jolly and Lakhotia, 2006; Prasanth et al., 2000; Singh and Lakhotia, 2015); (2) *flamenco*, which forms foci at the nuclear envelope in ovarian follicle cells and is important for egg chamber development and male courtship behavior (Mével-Ninio et al., 2007; Rivera et al., 2024; Sokolova et al., 2019); and (3) *CR34335* and *CR40469*, which are orthologs with roles in regeneration and development (Camilleri-Robles et al., 2024). The most abundant lncRNA was *CR45018*, which made up 41% of all lncRNAs in females and 30% in males (Figure 2a, b). However, this lncRNA has not been functionally characterized. Its only available annotation classifies it as a putative direct transcriptional target of the CREB/CRTC pathway, although this has not been validated (Wang et al., 2021). Although several of these lncRNAs have known functions, most remain uncharacterized, and even those with established roles may have additional, unexplored functions. The contact data generated by RD-SPRITE provide insight into the RNA and DNA environment surrounding specific lncRNAs, which can be used to generate testable hypotheses about lncRNA function and regulation in *Drosophila* in both sexes.

### 2.2 Several lncRNAs share RNA and DNA interactions with spliceosomal RNAs in males and females

Previous studies in humans have determined that lncRNAs such as MALAT1 may mediate alternative splicing activity by regulating the association of precursor messenger RNA (pre-mRNAs) and splicing factors (Huarte, 2015; Malakar, 2024, Ouyang et al. 2022; Miao et al. 2022). However, the regulation of alternative splicing by lncRNAs has not been explored in *Drosophila*. We surveyed lncRNA association with snRNAs by first identifying a subset of lncRNAs that share interactions with major snRNAs, U1, U2, U4, U5, and U6, and highlighted the top 10 (bolded) with the most snRNA interactors in our male and female RD-SPRITE clusters (Figure 2a). In both sexes, these highlighted lncRNAs are not the most abundant lncRNAs overall (Figure 2a), suggesting that snRNA co-occurrence may reflect a regulatory association rather than a byproduct of high lncRNA expression.

We selected the lncRNAs *CR45018*, *flamenco*, *CR45312*, *CR34335*, *CR40469*, and *Hsrω* to explore interactions with snRNAs because these are the top 6 snRNA interacting lncRNAs in females (Figure 2b). The lncRNA *CR40469* is the only lncRNA that is not also among the top 6 in males, but we analyzed the same lncRNAs in both sexes for a direct comparison (Figure 2b and 2c). To determine whether the selected lncRNAs share RNA interactions and bind to shared genomic loci with snRNAs, we generated RNA-RNA and RNA-DNA interaction profiles for all lncRNAs and the major snRNAs, U1, U2, U4, U5, and U6. On the RNA-RNA level, the lncRNA Hsrω which has known roles in splicing speckles (Prasanth et al., 2000; Jolly and Lakhotia 2006; Singh and Lakhotia 2015) interacts predominantly with snRNAs U1, U4, and U5 in both sexes (Figure 2b and 2c). On the RNA-DNA level, the lncRNA Hsrω binds shared genomic loci with snRNAs U1 and U2 in females (Figure 2e), and snRNAs U2, U4, and U5 in males (Figure 2f). The lncRNA *Hsrω* expressed in heat stress conditions, forms nuclear structures called omega speckles which are dynamic storage sites for splicing factors (Jolly & Lakhotia, 2006). Thus, the observed interactions between lncRNA Hsrω and splicing factors are consistent with previous studies, but its interaction with snRNAs is a novel finding.

### 2.3 LncRNAs *CR34335* and *CR40469* share RNA interactions and DNA interactions with snRNAs specifically in females

We also identified trends that are consistent with sex-biased differential interactions between a subset of lncRNAs and snRNAs (Figure 2d). Across snRNAs U1, U2, U4, U5, and U6, sex-biased enrichment was largely absent as expected for important spliceosomal RNAs (Figure 2d). The differential RNA-RNA interaction profile revealed that several lncRNAs show sex-biased enrichment with snRNAs. RNA pairs detected in only one sex (marked with a dot) were excluded from this comparison. Among pairs detected in both sexes, lncRNA *CR45018* was primarily enriched in males, whereas lncRNAs *CR40469* and *CR34335* were primarily enriched in females (Figure 2d).

We then explored whether lncRNAs organize within higher order structures with splicing factors, sharing multi-way interactions with snRNAs on chromatin (Figure 2g and 2h). We found that the lncRNAs *flamenco*, *CR45312*, and *CR45018* share multi-way interactions with each other, binding shared genomic loci in both females (Figure 2g; 84 genes) and males (Figure 2h; 33 genes), but do not simultaneously bind snRNAs. In contrast, the lncRNAs *CR34335* and *CR40469* show substantially more genomic overlap with snRNAs with a sex-biased pattern. A recent study that explored lncRNA function during regeneration identified lncRNA *CR40469* as an interesting target in development (Camilleri-Robles et al. 2024). Furthermore, lncRNA *CR34335* shares 99.1% sequence homology to lncRNA *CR40469* and was shown to partially compensate for lncRNA *CR40469* in function during regeneration (Camilleri-Robles et al., 2024). We observed more frequent associations of *CR34335* and *CR40469* with snRNAs in females (Figure 2g) than in males (Figure 2h). The female-biased chromatin association of lncRNA *CR34335* and *CR40469* with multiple snRNAs suggests that these lncRNAs may play a previously uncharacterized, sex-specific role in regulating alternative splicing.

Together, RNA-RNA and RNA-DNA interaction data reveal that several lncRNAs associate with spliceosomal components in a sex-biased manner, highlighting them as candidates for potential roles in sex-specific splicing.

### 3.1 CLAMP depletion alters snRNA-snRNA contacts and snRNA occupancy on chromatin

The pioneer transcription factor CLAMP (Chromatin-linked Adapter for MSL Proteins) is a key regulator of sex-specific alternative splicing in *Drosophila* males and females (Ray et al., 2023). CLAMP also binds several components of splicing machinery: 1) small nuclear RNAs (snRNAs), the core RNA components of the spliceosome (Ray et al., 2026) and 2) RNA-binding proteins (RBPs) such as Squid (Sqd), Hrp38, Hrp36, SC35 and Syncrip (Syp), which function in RNA processing, localization, and alternative splicing (Urban et al., 2017). Together, these findings suggest that CLAMP may reside within or associate with spliceosomal hubs. We therefore hypothesized that CLAMP mediates the organization of snRNA interaction within these hubs, both among snRNAs and between snRNAs and chromatin. Thus, we performed RD-SPRITE in male (S2) and female (Kc167) cell lines under control (GFP-RNAi) and established *clamp*-depleted (*clamp*-RNAi) conditions (Larschan et al. 2012) to reveal CLAMP-dependent snRNA-snRNA and snRNA-chromatin interaction effects (Table S1).

To explore the impact of CLAMP on snRNA organization, we compared RNA-RNA interaction profiles between GFP-RNAi control and *clamp*-RNAi cells in males and females (Figure 3a, 3b). We specifically focused on snRNA variants identified as direct CLAMP interactors by iCLIP (Ray et al., 2026; Figure S4a, S4b). As expected, we observed that variants of the same snRNA class interact with each other more often than variants from other classes (Figure 3a, 3b, S4c, S4e). Thus, in *clamp*-RNAi cells the overall interaction structure of snRNAs is preserved in both males and females as in GFP-RNAi samples. We then asked whether the frequency of snRNA contact pairs differed between conditions. In male cells, 16 pairs were more frequently identified in GFP-RNAi samples (log2FC > 1), 17 pairs were more frequently identified in *clamp*-RNAi samples (log2FC > 1), 13 pairs were only present in GFP-RNAi samples, 7 pairs were only present in *clamp*-RNAi samples, and 63 pairs were unchanged in frequency (Figure 3c). Most differential pairs involved snRNAs of different classes, not snRNAs of the same class (Figure 3a-d). In female cells, 2 pairs were higher in GFP-RNAi (log2FC > 1), 1 pair was higher in *clamp*-RNAi (log2FC > 1), 2 pairs were only present in GFP-RNAi, 5 pairs were only present in *clamp*-RNAi, and 7 pairs were unchanged (Figure 3d). Among contacts between snRNAs that are direct CLAMP targets that were detected in both GFP and *clamp* RNAi conditions, 34% changed after *clamp* RNAi in males (Figure 3c) and 30% in females (Figure 3d). Therefore, it is possible that CLAMP directly regulates the frequency of snRNA contacts within the major spliceosome complex because the snRNAs that directly interact with CLAMP and their contacts are altered by the depletion of CLAMP.

**Figure 3:**
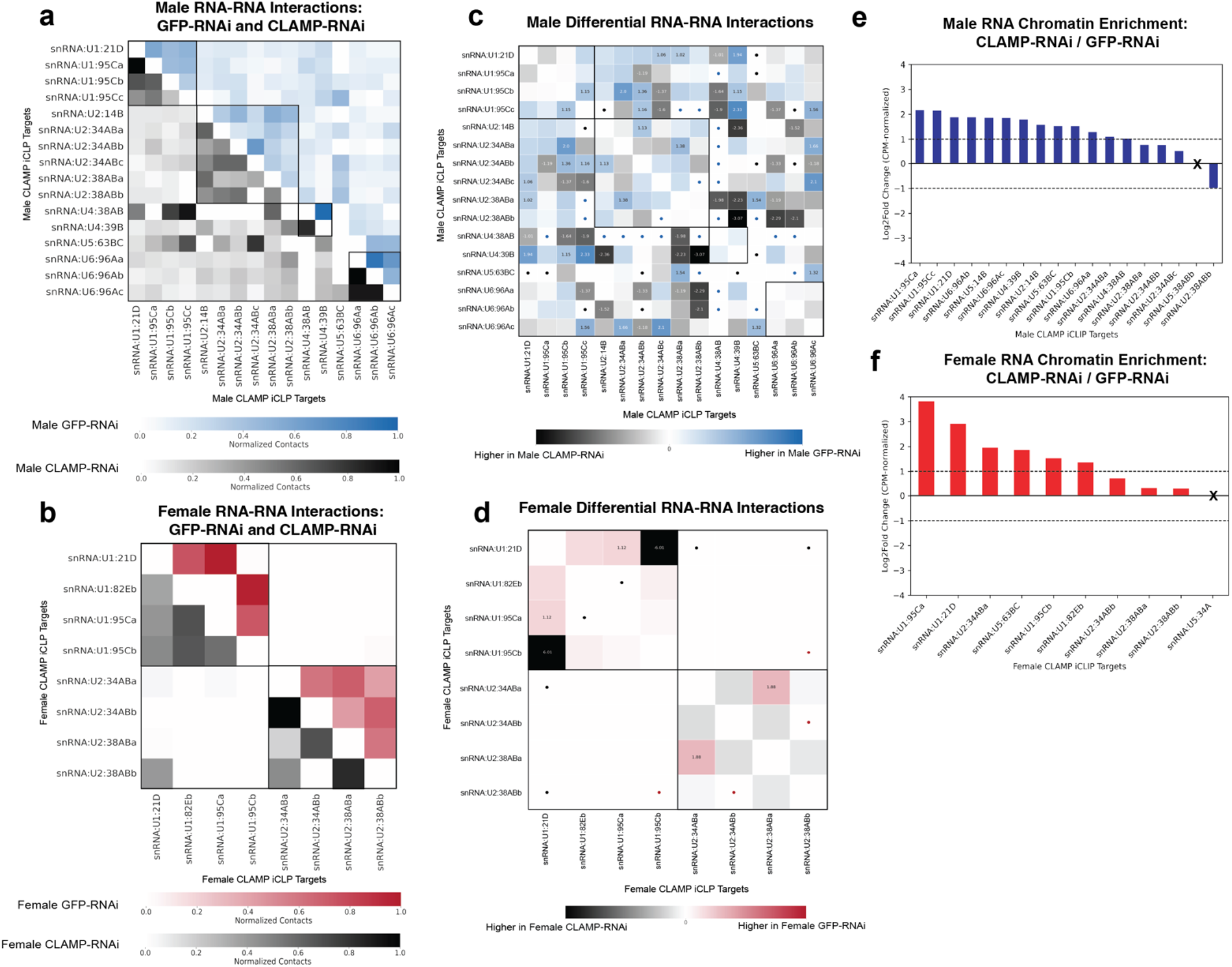
CLAMP depletion alters spliceosomal RNA interactions on the RNA and DNA level. All analyses in Figure 3 used the 10% male (S2) and female (Kc167) GFP-RNAi and CLAMP-RNAi libraries from Experiment B. **(a-b)** Heatmap showing pairwise RNA-RNA interactions between snRNA variants identified as direct CLAMP targets by iCLIP (Ray et al. 2026) in CLAMP-RNAi (bottom) and GFP-RNAi (top) cells. Boxes denote snRNA classes U1, U2, U4, U5, and U6. **(c-d)** Heatmap showing pairwise differential contacts calculated from male (c) and female (d) CLAMP-RNAi and GFP-RNAi RNA-RNA interaction profiles. Plotted RNAs are snRNAs identified as direct CLAMP targets by iCLIP (Ray et al. 2026). Normalized log2 fold change in RNA-RNA interaction frequency between GFP-RNAi and CLAMP-RNAi treated cells. For males, blue (right) indicates interactions higher in GFP-RNAi, and black (left) indicates interactions higher in CLAMP-RNAi. For females, red (right) indicates interactions higher in GFP-RNAi, and black (left) indicates interactions higher in CLAMP-RNAi. Values are shown for interactions with |log2 fold change| > 2. Dots mark interactions detected in either GFP-RNAi or CLAMP-RNAi, colored by the condition in which they were detected. **(e-f)** For each snRNA, weighted RNA-DNA contacts were normalized to counts per million (CPM) RNA-DNA clusters and summed across the genome. Bars show the log2 fold change of this value in CLAMP-RNAi relative to GFP-RNAi cells in male (e) and female (f) cells. The dotted line represents the 2-fold change threshold.

**Figure 4:**
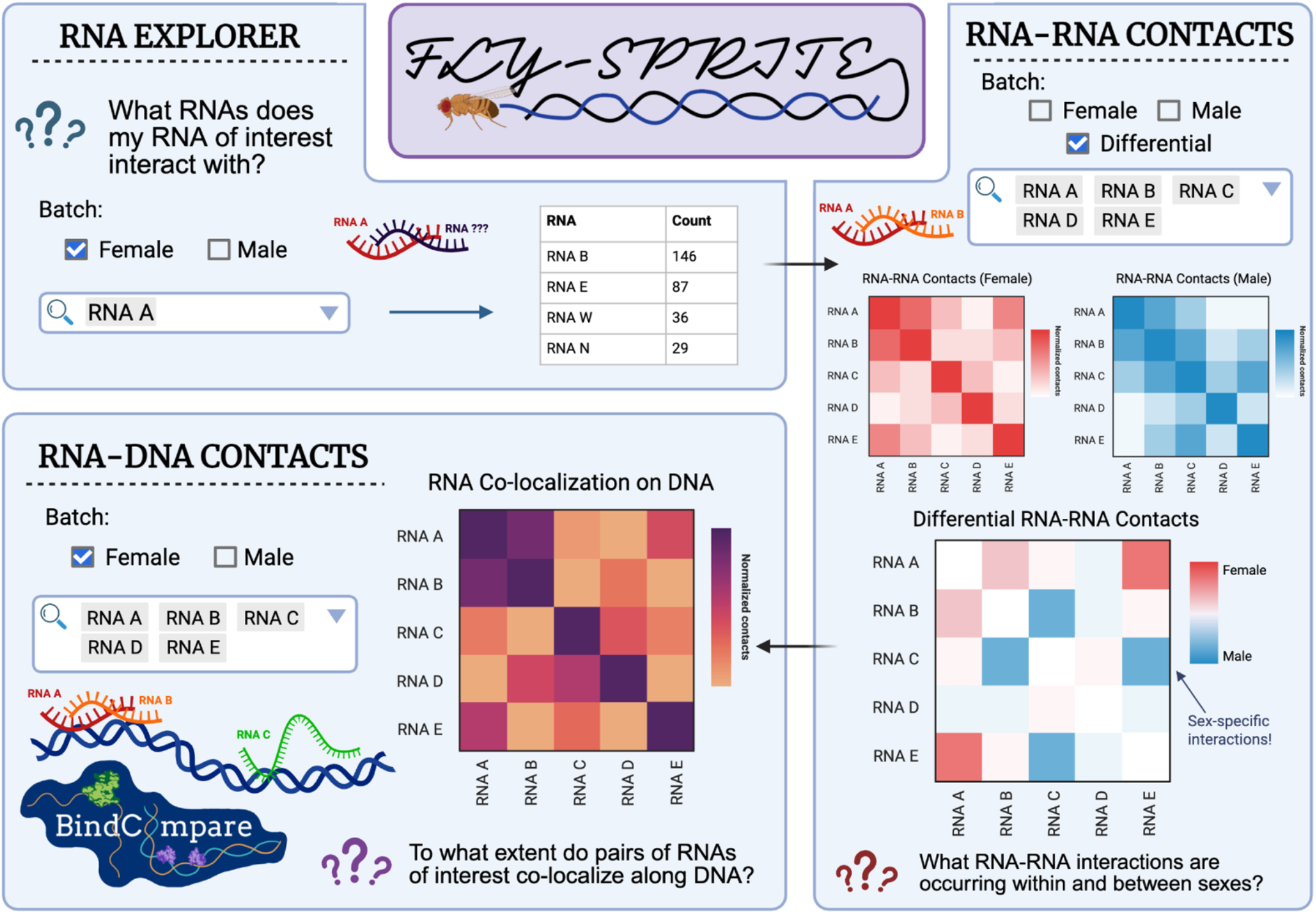
Schematic of the Fly-SPRITE platform. Users can interact with our *Drosophila* RD-SPRITE libraries through an interactive dashboard. Functions include exploration of candidate interactor RNAs with an RNA of interest, generation of RNA-RNA individual and differential contact matrices, and integration of BindCompare (Mahableshwarkar et al., 2024) to interrogate co-localization of RNAs along the genome.

To test whether *clamp* depletion impacts snRNA association with chromatin, we compared snRNA-DNA contacts between GFP-RNAi and *clamp*-RNAi cells in males and females. For each snRNA, we recorded RNA-DNA contacts genome-wide and normalized each contact to counts per million (CPM) RNA-DNA clusters in each library. We then summed all normalized values genome-wide and computed the log2 fold change between conditions (*clamp*-RNAi / GFP-RNAi) (Figure 3e, 3f, S4g, S4h). In both sexes, most snRNAs exhibited increased chromatin contacts in *clamp*-RNAi cells compared with GFP-RNAi cells. These results suggest that CLAMP depletion increases snRNA occupancy on chromatin, with the strongest effects on U1 snRNAs.

Together, these results suggest that CLAMP impacts snRNA organization at two levels: contacts between snRNAs and snRNA association with chromatin. Because both RNA-RNA interactions and chromatin association changed upon CLAMP depletion for RNAs that directly interact with CLAMP, it is possible that CLAMP prevents the formation of aberrant splice isoforms by regulating the composition of spliceosomal hubs and their targeting to chromatin.

### 4.1 The Fly-SPRITE tool makes *Drosophila* RD-SPRITE libraries accessible

We developed Fly-SPRITE, a dashboard created using Shiny (*Shiny for Python*) to facilitate broader community access to our *Drosophila* RD-SPRITE datasets. Fly-SPRITE is hosted as a web app, eliminating any need to download large datasets or maneuver complex pipelines. This tool will provide *Drosophila* researchers an accessible way to investigate the nuclear localization of individual RNAs or loci of interest without requiring a computational skillset.

We designed this tool specifically to work with our own RD-SPRITE libraries rather than accepting user-generated files due to the rarity of existing RD-SPRITE datasets and the resulting difficulty of ensuring consistency in file annotations between them. As RD-SPRITE becomes more widely used, Fly-SPRITE could be modified to work with user-inputted cluster files.

### 4.2 RNA-RNA Contacts

The first main tab of Fly-SPRITE allows the user to generate RNA-RNA contact matrices based on the methodology used to generate Figure 1b. First, a user can select which batches from our dataset they are interested in analyzing. QC information for each batch is available in Supplementary Table 2.

Once the batch(es) are selected, users can pick their RNAs of interest. A full list of RNAs with at least five reads in clusters in either one batch or in the union of both selected batches will populate based on the selected dataset. We also added larger categories of RNAs to the list of options in cases where we suspect that transcript-specific detail is less useful, such as all tRNAs grouped into a ‘tRNA (ALL)’ entry. Running the analysis generates normalized per-sample and differential heatmaps like those shown in Figures 1-3. Each individual RNA-RNA contact matrix in Fly-SPRITE is hierarchically clustered to indicate unbiased similarities between contact profiles of groups of RNAs.

### 4.3 RNA-DNA Contacts

The second main tab of Fly-SPRITE generates RNA-DNA contact matrices. Notably, this is the same type of analysis performed in Figure 1j, 2e, and 2f using BindCompare (Mahableshwarkar et al., 2024) which identifies shared chromatin targets of RNA pairs.

Once again, users are able to select a batch from our datasets and choose their RNAs of interest from the selection menu. This list of RNAs available for analysis in each batch is restricted to those found in clusters containing at least one DNA read. The script uses the bindexplore module of BindCompare (Mahableshwarkar et al., 2024) and returns correlation values between pairs of BED files, swapping the reference and overlapping files along the diagonal. This provides a measure of the degree to which two RNAs overlap in the location of their association with chromatin.

### 4.4 RNA Explorer Tool

In cases where a user wants to explore which RNAs interact with a particular RNA of interest, we provide an ‘RNA Explorer’ tab that returns top hits between a queried RNA and other RNAs within a batch. This can provide an idea of potential interactions to explore in more depth using the RNA-RNA or RNA-DNA tabs of Fly-SPRITE.

For example, a user can use Fly-SPRITE to investigate the function of a poorly characterized lncRNA, *CR45018*. If we use the RNA Explorer to determine which RNAs in our dataset it interacts with most frequently, we find that lncRNA:*flam* (*flamenco*) is the most frequent hit. Likewise, *CR45018* is the most common interactor when querying for *flam*. Observations like this can provide potential avenues for further exploration, either by utilizing Fly-SPRITE to find hubs of frequent interactors or RNAs that colocalize with each other on chromatin, or by performing downstream validation experiments to interrogate interaction structure and function.

## Discussion

The nucleus is home to multiple molecular compartments, or hubs, containing molecules with shared functional roles. These hubs are the sites of transcriptional activity, spliceosome function, ribosome biogenesis, and other processes, concentrating factors around genes to increase the frequency of binding events and improve the efficiency of gene regulation. The regulation occurring within these hubs is mediated by multiple ribonucleoprotein complexes, consisting of complex networks of proteins and RNA partners that are uniquely localized to the DNA of chromatin. Most molecular mapping techniques resolve pairwise interactions involved in gene regulation, including protein-RNA (iCLIP) (Huppertz et al. 2014), protein-DNA (ChIP-seq, CUT&RUN) (Skene and Henikoff 2017; Solomon et al. 1988), RNA-DNA (ChAR-seq) (Bell et al. 2018), and DNA-DNA interactions (Hi-C, Micro-C) interactions (Belton et al. 2012; Hsieh et al. 2015). Higher-order interactions among RNAs, DNAs, and proteins within the same complex or hub can sometimes be inferred by combining these assays, but this approach requires many experiments and is costly and time-consuming. Therefore, we implemented, for the first time, RNA-DNA Split Pool Recognition of Interactions by Tag Extension (RD-SPRITE) (Quinodoz et al. 2021) in *Drosophila* male (S2) and female (Kc167) cells to identify all multi-way nucleic acid interactions within ribonucleoprotein nuclear hubs genome-wide and characterize any potential sex-specific trends.

Using RD-SPRITE, we identified RNA hubs containing nucleolar, spliceosomal, and cytoplasmic RNAs (Figure 1). RNAs within each group frequently occurred in the same clusters and also contacted similar genomic regions. These regions were enriched in transcriptionally active chromatin, where their RNA processing complexes function. Hub formation by nucleolar, spliceosomal, and cytoplasmic RNAs was consistent with previous observations in human cells (Quinodoz et al., 2021). However, RNAs in *Drosophila* hubs more frequently interacted with RNAs from other functional classes than those in mammals. One explanation may be differences in nuclear organization. The *Drosophila* genome is about 5% the size of the human genome, yet the two encode a comparable number of genes, about 15,500 across four chromosome pairs in *Drosophila* and about 22,000 across 23 chromosome pairs in humans (Adams et al., 2000; Aquadro et al., 2001; Nurk et al., 2022). *Drosophila* nuclei are also smaller, at about 78 µm³ compared with 370-690 µm³ in human cells (Maul et al., 1977). As a result, *Drosophila* nuclei contain 3-6x more genes per unit volume (*Drosophila*: 198 genes/µm^3, humans: 32-59 genes/µm^3). This higher gene density may limit the spatial segregation of RNAs by functional class, allowing RNAs from different functional groups to occupy shared nuclear space. Furthermore, spliceosomal and nucleolar hub components are highly conserved across species suggesting that the similar hub components exist but are present within a much smaller nuclear volume.

To characterize interactions among spliceosomal RNAs, we identified higher-order contacts among snRNAs and other RNAs that are potential components of spliceosomal hubs. Several lncRNAs co-occured with snRNAs in both male (S2) and female (Kc167) cells, but their interaction patterns differed. One notable candidate was the antisense RNA, asRNA *CR45330*, which interacts more with the U2 snRNAs than other snRNAs classes in both sexes (Figure 1). During spliceosome assembly, U2 snRNA and snRNP base-pairs with the intron branch point sequence, positioning the branch point for the first catalytic step of splicing (Ares and Weiser 1995). These data suggest that asRNA *CR45330* may function with U2 during branch point recognition. However, co-occurrence within a hub does not establish a functional role. Future work will be needed to validate asRNA *CR45330* interactions with U2 spliceosomal components and test its potential regulatory role in U2-intron splicing.

We also identified two X-linked lncRNAs, CR34335 and CR40469, whose co-occurrence with snRNAs was female-biased (Figure 2). These lncRNAs share high sequence similarity and are essential to *Drosophila* development (Camilleri-Robles et al., 2024), but their role in splicing has not been explored. X-linked genes often show sex-biased expression and regulation, shaped by dosage compensation and other X-specific evolutionary pressures (Parisi et al., 2003; Meiklejohn et al., 2011; Meiklejohn and Presgraves, 2012). This chromosomal context may contribute to the sex-biased associations we observed. However, differences in lncRNA expression, or other differences between S2 and Kc167 cells, may also contribute. The interaction of *CR34335* and *CR40469* with snRNAs suggests that they may play a previously uncharacterized role in AS, possibly including sex-specific AS. Depleting these lncRNAs in male and female cells and measuring changes in AS would test whether these lncRNAs regulate splicing, and whether their effects are sex-specific.

CLAMP (Chromatin-Linked Adaptor for MSL Proteins) is a *Drosophila* DNA-binding protein that regulates sex-specific alternative splicing (AS) in both males and females (Ray et al. 2023). CLAMP depletion impacts sex-specific splicing activity genome-wide, producing aberrant isoforms and increasing cryptic splicing (Ray et al. 2023). However, how CLAMP influences splicing regulation remains unclear. In particular, it is unknown how CLAMP affects interactions among snRNAs and the organization of spliceosomal hubs. To address this gap, we used RD-SPRITE to profile the RNA-RNA and RNA-DNA interactions of CLAMP-bound snRNAs, identified by iCLIP (Ray et al. 2026), in male (S2) and female (Kc167) cells. We then compared these interactions between control and *clamp* RNAi conditions.

The overall organization of snRNA-snRNA contacts was largely maintained after CLAMP depletion in both sexes, with snRNAs continuing to group by class (Figure 3). In contrast, snRNA association with chromatin increased after CLAMP depletion in both male and female cells (Figure 3). These results suggest that CLAMP may restrict snRNA-chromatin association. One model is that CLAMP limits spliceosomal engagement at inappropriate sites, which would be consistent with its role in suppressing cryptic sex-specific splicing. Together, our data support a role for CLAMP in organizing spliceosomal RNAs on chromatin. It also identifies specific snRNA within the spliceosome as candidates for studying how CLAMP-dependent RNA interactions contribute to AS and the suppression of cryptic splicing, which would require further validation.

The nucleus controls the developmental program and how cells respond to internal and external changes (Shan et al., 2024). Beyond the hubs and lncRNAs described in this study, RD-SPRITE also captures other regulatory RNAs of both high and low abundance. These include a broad range of RNAs, including miRNAs, snoRNAs, scaRNAs, transcripts of many protein-coding genes, and numerous less-characterized non-coding RNAs. Our RD-SPRITE *Drosophila* male (S2) and female (Kc167) data, for the first time, capture these RNA simultaneously and their interaction with other RNA partners and specific DNA regions on chromatin. Having integrated our datasets and analysis scripts into Fly-SPRITE, researchers can now explore the RNA and DNA contacts of RNAs relevant to their work in male S2 and female Kc167 cells, including candidate sex-biased RNA-RNA contacts.

Using RD-SPRITE in *Drosophila* cells, we: (1) identified RNA hubs containing nucleolar, spliceosomal, and cytoplasmic RNAs some of which show interactions with each other not observed in mammalian cells, (2) identified lncRNAs that interact with snRNAs, some with sex-dependent specificity, (3) found that CLAMP depletion increases the association of specific snRNAs with chromatin, and (4) developed Fly-SPRITE, a platform for exploring RNA-RNA and RNA-DNA interactions of any RNA of interest in male and female cells.

## Methods and Materials

### 1. Experimental Methods

#### 1.1 Cell culture

Kc167 and S2 cells were maintained at 25°C in Schneider’s media supplemented with 10% Fetal Bovine Serum and 1.4X Antibiotic-Antimycotic (Thermofisher Scientific, USA). Cells were passed every three days to maintain an appropriate cell density.

#### 1.2 RD-SPRITE library preparation

Cell lysate was prepared and DNA fragmentation performed according to the SPRITE protocol (Quinodoz et al 2022). Standardization for the MNAse concentration used for digestion was performed and a 1:500 MNase dilution was used. Lysate coupling to NHS beads and RD-SPRITE library preparation after the split-pool barcoding procedure was done according to the RD-SPRITE protocol in Quinodoz et al 2018. 1.5 ml NHS beads per sample was used at a bead: molecule ratio of 2. Two replicates of Kc167 and S2 cells treated with *clamp* dsRNA and GFP dsRNA *clamp*-RNAi and GFP-RNAi were used (Aguilera et al. 2026). For one set of Kc167 and S2 GFP-RNAi samples, a non-modified DPM tag was used. For the rest of the samples, a biotin-modified DPM tag was used. The split-pool barcoding procedure was performed according to Quinodoz et al., (2022). Five rounds of split-pool barcoding, including DPM tag, RPM tag, ODD tag, EVEN tag, ODD tag and finally Terminal tag. 2.5%, 5%, and 10% aliquots from each sample were used for library preparations. Each library was barcoded with a different adaptor according to Quinodoz et al. 2018.

### 2. Computational Methods

#### 2.1 RD-SPRITE computational pipeline details and adaptations for Drosophila genomes

We adapted the RD-SPRITE analysis pipeline from Quinodoz et al. (2021) (https://github.com/GuttmanLab/sprite2.0-pipeline) for *Drosophila melanogaster*. All alignments used the dm6 genome assembly and FlyBase annotation release 6.60.

##### Sequencing and quality control

Libraries were sequenced on Novaseq platform using 150x pair-end sequencing. We trimmed adapters from raw reads with Trim Galore v0.6.2 (citation). We assessed read quality with FastQC v0.11.8 (citation). We summarized QC reports with MultiQC v1.6 (citation).

##### Barcode identification

We identified the split-pool barcode on each read pair with Barcode ID v1.2.0.jar (citation). We kept only reads that carried all five tags. Reads with incomplete barcodes cannot be assigned to a single cluster, so we discarded them. Ligation efficiency for each round is reported in Supplementary Table 3 (insert here).

##### RNA and DNA read separation

We split barcoded reads into RNA and DNA files based on the adapter on Read 1. RNA reads carry the RPM adaptor and DNA reads carry the DPM adaptor. We then trimmed the RPM and DPM sequences from the 5’ end of Read 1, and any barcode sequence from the 3’ end, with Cutadapt v2.5 (citation).

##### RNA alignment

We aligned RNA reads in two steps. First, we aligned RNA reads to a repeat-aware index of highly abundant noncoding RNAs with Bowtie v2.3.5 (citation). This index groups multi-copy RNAs, such as rRNAs, tRNAs, and other non-coding RNAs into broad categories based on their FlyBase IDs. Second, we aligned the remaining reads to dm6 with the splice-aware aligner HISAT2 v2.1.0 (citation). We assigned reads to genes with featureCounts (Subread v1.6.4) (citation) using the FlyBase GTF (6.60).

##### DNA alignment

We aligned DNA reads to dm6 with Bowtie v2.3.5 (citation). We removed reads with overlapping dm6 blacklist regions (Amemiya et al. 2019).

##### Cluster assembly and filtering

We grouped all RNA and DNA reads that shared an identical barcode into a single cluster. Following Quinodoz et al. 2021, we kept clusters containing 2 to 1,000 reads. Refer to Supplementary Figure 1 for information on cluster size distributions for Kc167 (female) and S2 (male) batches.

### 2.2 Analysis of RNA-RNA interactions

#### 2.2.1 RNA-RNA interaction matrices

For each cluster, we identified the unique RNAs present and counted each pair of distinct RNAs once. We summed these counts across clusters to build a symmetric RNA-RNA contact matrix. Each entry is the number of clusters containing both RNAs. For each cluster, we identified the unique RNAs present and counted each pair of distinct RNAs once. We summed these counts across clusters to build a symmetric RNA-RNA contact matrix. Each entry is the number of clusters containing both RNAs. We set self-interactions to zero and normalized each RNA-RNA contact matrix with iterative correlation and eigenvector decomposition (ICE; Imakaev et al. 2012), as in Quinodoz et al. (2021). We used the numutils.iterative_correction_symmetric function from the Python package cooltools (v0.7.1) with default parameters. ICE is a matrix-balancing method that corrects for biases in contact coverage that arise from unequal RNA abundance. However, we found that RNAs with low read counts can produce inflated values after ICE normalization. We therefore removed RNAs with fewer than 5 total reads. To choose this cutoff, we computed the variance of the ICE-normalized matrix across minimum read cutoffs from 0 to 20 (Supplementary Figure 6). The variance was stable at cutoffs of 5 and above in both male (S2) and female (Kc167) RD-SPRITE batches. We applied the same RNA abundance cutoff to all RNA-RNA interaction matrices prior to normalization.

#### 2.2.2 Multi-way Contact Score

The multi-way contact score tests whether a set of k-mer RNAs co-occurs in clusters (observed) more often than expected by chance. We reimplemented this score following Quinodoz et al. 2021. For a k-mer interaction such as A-B-C, we first selected clusters containing one (k-1) RNA sub-fragment, such as A-B. In each selected cluster, we replaced the remaining RNAs with random draws. Each RNA was drawn with a probability equal to its fractional abundance – defined as (# RNAs A / total # RNAs). We then computed the clusters that contained the full k-mer in the permuted dataset. We repeated this for each of the k-mer sub-fragments (A-B, A-C, B-C) for 100 iterations to generate an expected distribution. For each k-mer sub-fragment, we computed a p-value and z-score from the observed count and the expected distribution over 100 iterations. Consistent with Quinodoz et al. 2021, we reported the maximum statistics of all possible paths to assembling the k-mer (e.g., max(A-B|C, B-C|A, A-C|B)).

#### 2.2.3 Differential RNA-RNA interaction matrices

We compared ICE-normalized matrices following the framework of diffHiC (Lun & Smyth 2015). Each sample contributes a library-specific bias from uncontrolled differences in library preparation. Such technical differences may manifest as trended differences between libraries, where the magnitude of the difference varies as a function of the average read abundance. To remove trended biases, we kept only RNA pairs present in both matrices. When at least 20 RNA pairs were shared, we then applied cyclic loess normalization adapted for low counts (Lun & Smyth 2016), as suggested by diffHiC (Lun & Smyth 2015). We fit LOESS curves using the loess function from the Python package scikit-misc (v0.3.1). Cyclic loess is an iterative normalization method that repeatedly applies loess (locally weighted regression) smoothing to pairwise MA (minus and average) plots of samples until the data converges, removing intensity-dependent biases across samples. When fewer than 20 RNA pairs were shared, a loess fit was not reliable, so we applied mean scaling instead. We adjusted each comparison so that the mean log2 ratio across shared pairs was zero. This removes the overall difference in library scale but not intensity-dependent trends. For each shared RNA pair, we computed a log2 fold change. Positive values indicate enrichment in sample 1. Negative values indicate enrichment in sample 2. Because we had one biological replicate per condition, we report log2 FC values descriptively. We considered pairs with |log2 FC| ≥ 1 candidate differential interactions.

### 2.3 Analysis of RNA-DNA interactions

#### 2.3.1 Generating RNA-DNA contact profiles

We measured genome-wide localization for each RNA by recording the 1kb genomic bins in which the RNA co-occured. We used a python script, rna_dna_contacts.py, from Quinodoz et al. 2021 on the male (S2) and female (Kc167) cluster files. We analyzed the major dm6 chromosome arms: chr2L, chr2R, chr3L, chr3R, chr4, and chrX. We included chrY for S2 cells only. For each RNA, we exported the contacted bins as both BED files and bedgraphs. For the bedgraphs, consistent with Quinodoz et al. 2021, we weighted each RNA-DNA contact by 2/*n*, where *n* is the number of reads in the RNA’s corresponding cluster.

#### 2.3.2 Chromatin state analysis of RNA and DNA contacts

To identify chromatin environments that nucleolar and spliceosomal RNAs interact, we compared RNA-DNA contacts with published chromatin state models. For S2 cells, we used the 9-state HMM model from Kharchenko et al. (2011), and for Kc167 cells, we used the 5-state HMM model from Filion et al. (2010). We divided the dm6 genome into 1kb bins and assigned each bin its overlapping chromatin states using bedtools map (v2.27.1). Genomic bins that overlapped more than one state were annotated all overlapping states. Bins with no overlapping state were excluded.

For each RNA, we identified the 1kb genomic bins it contacts from our RNA-DNA RD-SPRITE clusters in a BED file as described in 2.3.1. We then combined bins across RNAs within each broad biological group to define contact bins for nucleolar RNAs (U3 snoRNAs, snoRNAs excluding U3 snoRNAs, ETS, ITS1, and ITS2) and spliceosomal RNAs (snRNA U1, U2, U4, U5, and U6). For each chromatin state, we calculated the fraction of RNA-DNA contacts in that state and the fraction of all genomic bins in that state. Bins with more than one chromatin state contributed to each of their states in both calculations. Fold enrichment was calculated as the fraction of RNA-DNA contacts in a state divided by the fraction of genomic bins in that state. Values above 1 indicate more contacts than expected from the genomic coverage of that state. We also repeated this analysis for all RNA-DNA contacts in our RD-SPRITE data to determine the overall distribution of chromatin states contacted by RNA.

#### 2.3.3 Pairwise co-localization of RNA-DNA contacts

We identified genomic regions where distinct RNA transcripts co-localize on the genome using the BindExplore module of BindCompare (Mahableshwarkar et al., 2024). BindExplore intersects the BED files of each RNA pair. For each pair, BindExplore reports an overlap fraction: (# bins contacts by both A and B) / (# bins contacts by reference RNA). This score is asymmetric because it depends on which RNA is the reference. We then identified genes contacted by each RNA and pairs of RNAs with BindCompare and the Flybase GTF (6.60). Genes were assigned if any region of its gene body overlaps within the 1kb genomic bin.

#### 2.3.4 Multi-way co-localization of RNA-DNA contacts

We tested whether genes contacted by two or more RNAs overlap more than expected by chance. We used the SuperExactTest (Wang et al. 2015) on the gene sets from 2.3.3. The SuperExactTest enumerates the elements shared by every possible combination of the sets and then computes fold change and the one-sided probability for assessing statistical significance represented as p-values of each observed intersection (Wang et al., 2015). The background population was 17200 genes, the total number of genes in the FlyBase GTF (6.60). We corrected p-values with Benjamini-Hochberg and used a threshold of FDR < 0.05. We treated significant intersections as candidate multi-RNA hubs on chromatin.

## Data and code availability

RD-SPRITE datasets for the Kc167 (female) and S2 (male) control and CLAMP-RNAi conditions generated during this study have been deposited on GEO and are publicly available as of the date of publication at [GEO link pending]. Code for the Fly-SPRITE application is available at [GitHub link pending]. This repository also includes the RNA-RNA and RNA-DNA analysis scripts, adapted from Quinodoz et al 2021, with documentation for their use.

## Acknowledgments

This work was supported by R35GM126994 to E.N.L. from NIH. S.G. was a trainee supported under the Brown University Predoctoral Training Program in Biological Data Science (NIH T32 GM149433). We thank Sofia Quindoza and Mitch Guttman for technical assistance with the RD-SPRITE protocol. We thank Mario R. Blanco for helping with computational analysis using the RD-SPRITE pipeline established by the Guttman Lab.

## Author Contributions

**S.G**. and **E. N. L.** planned experiments, analyzed results, and wrote the manuscript. **M.R.** planned the experiments, analyzed results, reviewed the manuscript, conducted the experimental work, and collected data for RD-SPRITE. **S.G**. performed all the computational and statistical analyses and established the RD-SPRITE pipeline for *Drosophila* data analysis. **M.C.** helped create the *Drosophila* genomic data-compatible RD-SPRITE pipeline and created the Fly-SPRITE platform.

## Supplementary Materials

**Supplementary Table 1.** RD-SPRITE library statistics for samples in Experiment A and Experiment B. Experiment A includes untreated control libraries. Experiment B includes GFP-RNAi control libraries and CLAMP-RNAi libraries from two biological replicates (BR1 and BR2). For each condition, male (S2) and female (Kc167) libraries were prepared from a single sample at two dilutions: 2.5% and 5% in Experiment A, and 5% and 10% in Experiment B. Ligation efficiency is the estimated per-round efficiency of barcode ligation. It was calculated as *f* ^ (1/5), where *f* is the fraction of reads containing all five barcode tags. Values were averaged across technical replicates. RPM and DPM reads are RNA and DNA reads, respectively. Cluster counts include only clusters containing 2-1,000 reads. Refer to cluster size distributions for each library in Supplementary Figure 1. RNA-DNA clusters contain at least one RNA read and at least one DNA read. RNA-DNA clusters were used for RNA-DNA interaction analyses. RNA and RNA-DNA clusters contain at least one RNA read, with or without DNA reads. RNA and RNA-DNA clusters were used for RNA-RNA interaction analyses.

| Experiment A – Control | Male Control (2.5%) | Female Control (2.5%) | Male Control (5%) | Female Control (5%) |
|---|---|---|---|---|
| Ligation efficiency each round (averaged across technical replicates) | 81% | 73% | 82% | 66% |
| Filtered # Reads | 920,355 | 598,848 | 501,266 | 403,852 |
| Filtered # RPM Reads | 896,779 | 579,829 | 477,928 | 389,397 |
| Filtered # DPM Reads | 23,576 | 19,019 | 23,338 | 14,455 |
| # Clusters (2-1000) | 168,107 | 114,639 | 128,269 | 90,543 |
| # RNA-DNA Clusters (2-1000) | 1,107 | 849 | 1,533 | 1,940 |
| # RNA and RNA-DNA Clusters (2-1000) | 164,250 | 111,672 | 26,447 | 12,650 |
| **Experiment B – GFP-RNAi** | **Male GFP-RNAi (5%)** | **Female GFP-RNAi (5%)** | **Male GFP-RNAi (10%)** | **Female GFP-RNAi (10%)** |
| Ligation efficiency each round (averaged across technical replicates) | 65% | 59% | 67% | 62% |
| Filtered # Reads | 146,001 | 107,979 | 281,747 | 96,407 |
| Filtered # RPM Reads | 143,105 | 94,122 | 274,071 | 80,594 |
| Filtered # DPM Reads | 2,896 | 13,857 | 7,676 | 15,813 |
| # Clusters (2-1000) | 21,409 | 21,659 | 27,204 | 15,261 |
| # RNA-DNA Clusters (2-1000) | 721 | 1,618 | 1,533 | 1,940 |
| # RNA and RNA-DNA Clusters (2-1000) | 21,039 | 19,747 | 26,447 | 12,650 |
| **Experiment B – CLAMP-RNAi Biological Replicate (BR) 1** | **Male CLAMP-RNAi (5%)** | **Female CLAMP-RNAi (5%)** | **Male CLAMP-RNAi (10%)** | **Female CLAMP-RNAi (10%)** |
| Ligation efficiency each round (averaged across technical replicates) | 71% | 66% | 70% | 70% |
| Filtered # Reads | 63,823 | 170,161 | 298,792 | 113,318 |
| Filtered # RPM Reads | 49,929 | 148,337 | 210,711 | 88,780 |
| Filtered # DPM Reads | 13,894 | 21,824 | 88,081 | 24,538 |
| # Clusters (2-1000) | 11,509 | 54,994 | 36,481 | 37,139 |
| # RNA-DNA Clusters (2-1000) | 1,092 | 1,256 | 4,421 | 752 |
| # RNA and RNA-DNA Clusters (2-1000) | 10,240 | 50,944 | 28,440 | 30,640 |
| **Experiment B – CLAMP-RNAi Biological Replicate (BR) 2** | **Male CLAMP-RNAi (5%)** | **Female CLAMP-RNAi (5%)** | **Male CLAMP-RNAi (10%)** | **Female CLAMP-RNAi (10%)** |
| Ligation efficiency each round (averaged across technical replicates) | 71% | 57% | 83% | 66% |
| Filtered # Reads | 409,587 | 108,606 | 643,639 | 117,750 |
| Filtered # RPM Reads | 409,339 | 108,406 | 642,883 | 117,673 |
| Filtered # DPM Reads | 248 | 200 | 756 | 77 |
| # Clusters (2-1000) | 123,961 | 36,572 | 119,680 | 26,287 |
| # RNA-DNA Clusters (2-1000) | 149 | 89 | 371 | 48 |
| # RNA and RNA-DNA Clusters (2-1000) | 123,951 | 36,543 | 119,654 | 26,281 |

### Note

Library quality, including ligation efficiency, RPM and DPM read counts, and cluster counts affected our ability to analyze pairwise and multi-way RNA-RNA and RNA-DNA interactions. Because this study compares sexes, we paired the highest quality male and female libraries from the same experiment and dilution. This controls for differences introduced by dilution. For control analyses (Fig. 1, Fig. 2, and Supplementary Fig. 1), we used the 2.5% male and female libraries from Experiment A.

CLAMP-RNAi and GFP-RNAi libraries also differed in RPM and DPM read depth. BR1 libraries were suitable for both RNA-RNA and RNA-DNA analyses. For Fig. 3, we used the 10% GFP-RNAi and CLAMP-RNAi BR1 male and female libraries. BR2 libraries had too few DNA reads for RNA-DNA analyses but were suitable for RNA-RNA analyses. For Supplementary Fig. 3, we used the male and female 10% BR2 for RNA-RNA analyses and the male and female 5% BR1 for RNA-DNA analyses. The male and female 5% BR2 libraries were not used for analyses but are publicly available.

All Experiment A control and Experiment B GFP-RNAi datasets, including both sexes and all dilutions, are available in Fly-SPRITE.

---

**Supplementary Figure 1.**
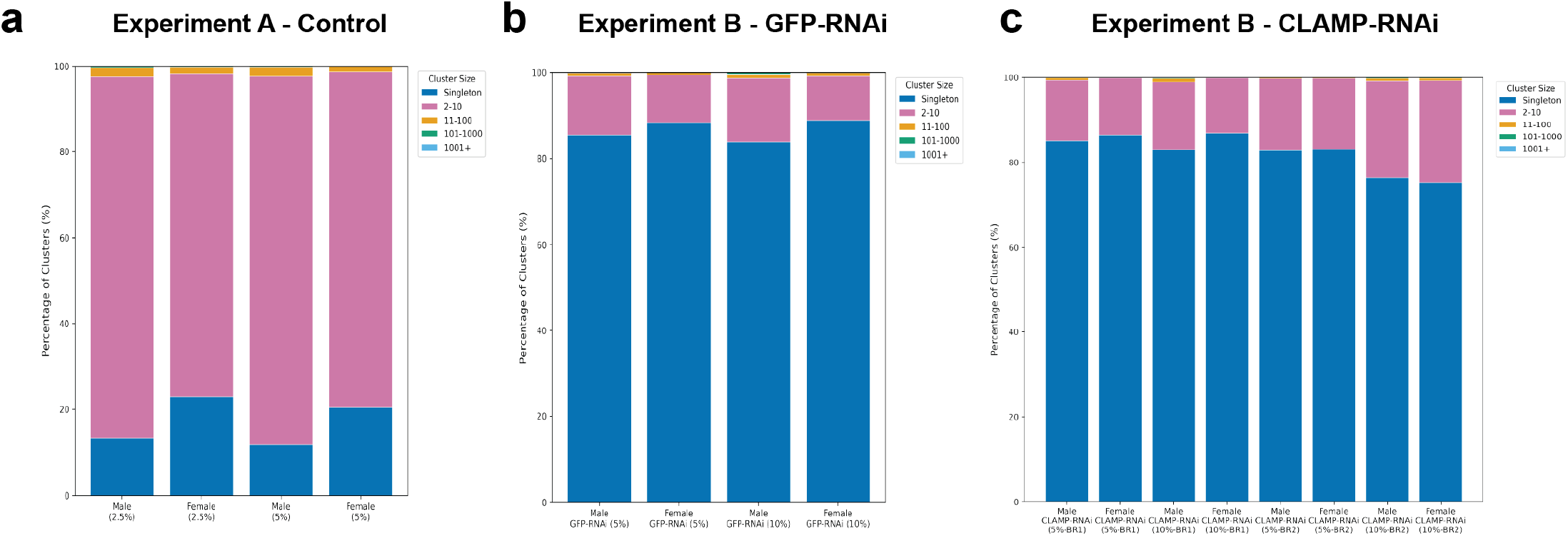
Cluster size distributions across RD-SPRITE libraries. Percentage of clusters in each size category (singleton, 2-10, 11-100, 101-1,000, and more than 1,000 reads) for male (S2) and female (Kc167) libraries. (a) Experiment A control libraries at 2.5% and 5% dilutions. Clusters of 2-10 reads were most common (75.4-85.7%), followed by singletons (11.9-22.8%), clusters of 11-100 reads (1.3-2.1%), 101-1,000 reads (0.04-0.23%). And more than 1,000 reads (0-0.07%). (b) Experiment B GFP-RNAi libraries at 5% and 10% dilutions. Singletons were most common (83.8-88.9%), followed by clusters of 2-10 reads (10.3-14.9%), 11-100 reads (0.46-0.88%). 101-1,1000 reads (0.07-0.36%) and more than 1,000 reads (0.006-0.07%). (c) Experiment B CLAMP-RNAi libraries from biological replicates 1 and 2 at 5% and 10% dilutions. Singletons were most common (75.1-86.8%), followed by clusters of 2-10 reads (13.1-24.1%), 11-100 reads (0.08-0.94%), 101-1,000 reads (0.02-0.22%), and more than 1,000 reads (0.006-0.07%). Only clusters containing 2-1,000 reads were used for downstream analyses.

**Supplementary Figure 2.**
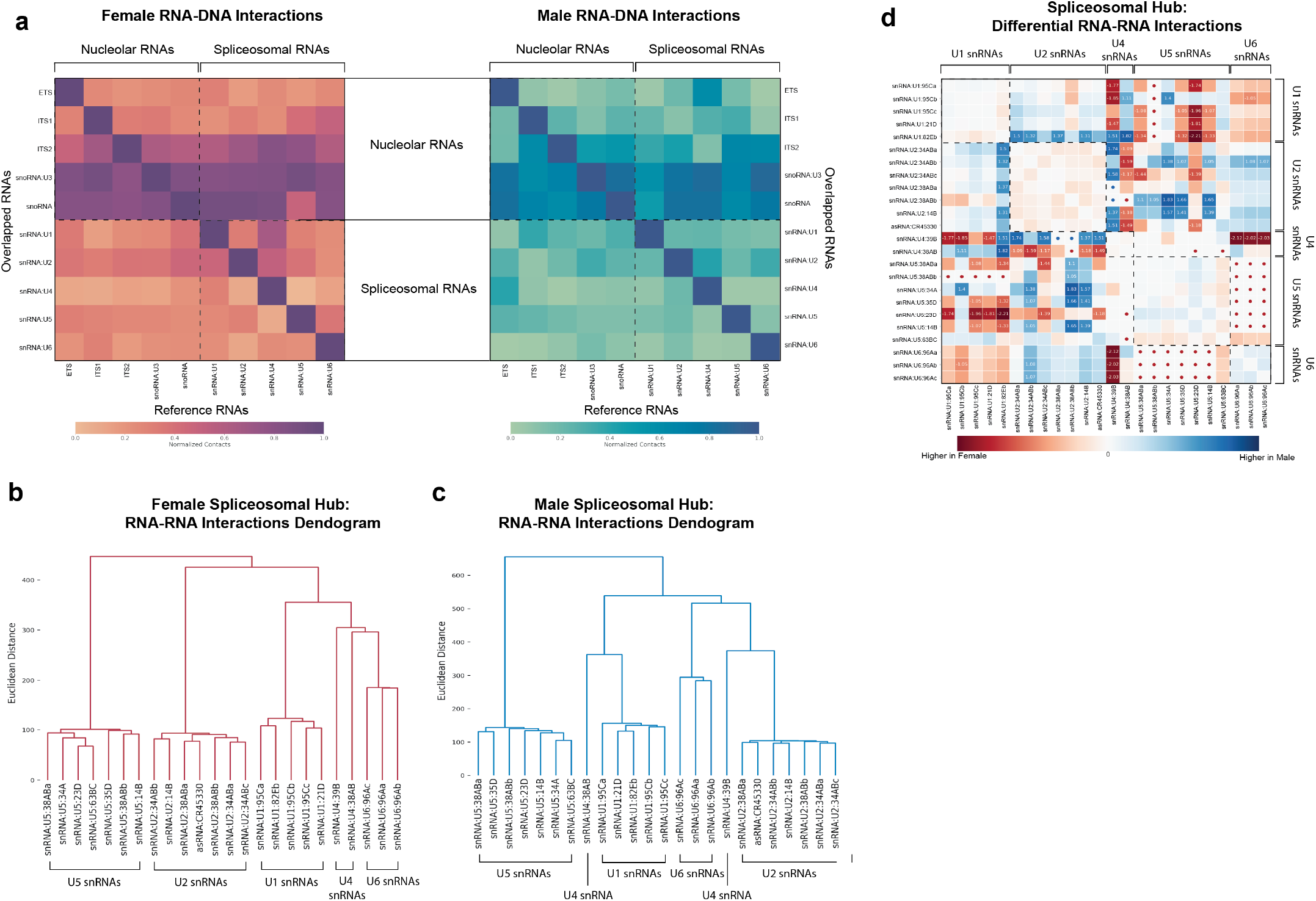
RD-SPRITE measures hub formation among RNA and DNA contacts in male and female cells, related to. Figure 1. All analyses in Supplementary Figure 2 used the 2.5% male (S2) and female (Kc167) control libraries from Experiment A. **(a)** Heatmap showing overlapped RNA-DNA binding sites between RNAs involved in nucleolar and spliceosomal function in female (left) and male (right) cells. The purple/orange and blue/green for females and males, respectively, represent high and low correlations of pairwise RNA localization to DNA. Boxes denote functional groups of RNAs. **(b)** Hierarchical clustering of snRNAs based on their RNA-RNA interaction profiles in female cells. Distances are Euclidian with ward linkage. snRNAs are grouped by class (U1, U2, U4, U5, and U6) with lncRNA CR45330. **(c)** Same analysis as in (b), for males. snRNAs again are grouped by class, although the arrangement of class clusters slightly differed from female cells. **(d)** Normalized log2 fold change in RNA-RNA interaction frequency between male and female cells. Red indicates interactions higher in female cells, and blue indicates interaction higher in male fells. Values are shown for interactions with |log2 fold change| > 2. Dots mark interactions detected in only one sex, colored by the sex in which they were detected. Lines separate snRNA classes. The lncRNA CR45330 is plotted with U2 snRNAs.

**Supplementary Figure 3.**
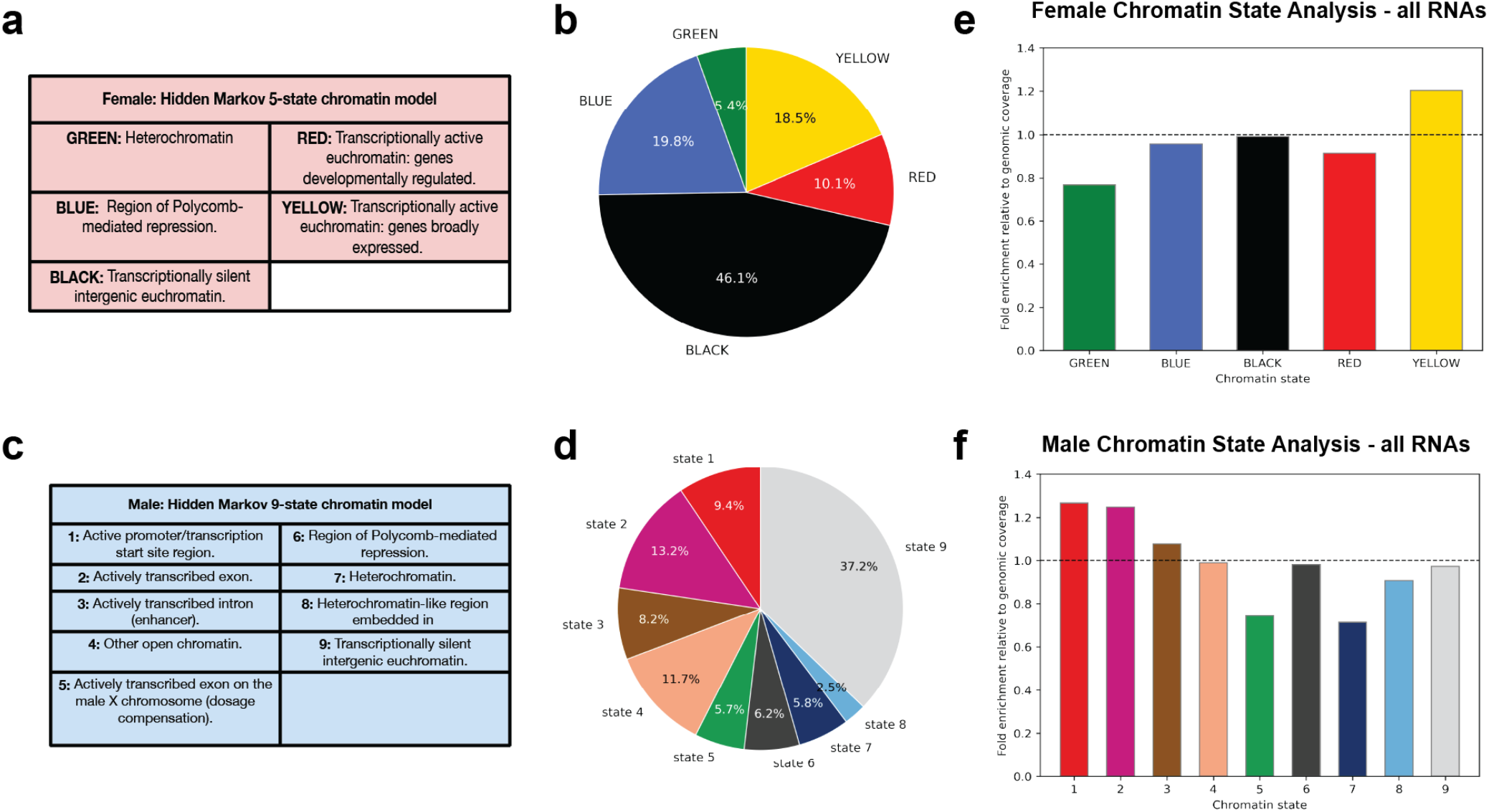
Genome-wide distribution of chromatin states in female and male cells. Percent of the genome assigned to each chromatin state. **(a-b)** Female (Kc167) cells, annotated with a five-state hidden Markov model (Fillion et al., 2010). **(c-d)** Male (S2) cells, annotated with a nine-state hidden Markov model (Kharchenko et al., 2011). The two models use different states, so proportions are not directly comparable between sexes. **(e-f)** For each chromatin state, the fraction of RNA-DNA contacts falling in that state, normalized to the fraction of the genome it covers. All RNAs in 2.5% control clusters from female (e) and male (f) cells, Experiment A. The dashed line marks the expected value of 1. Values above 1 indicate enrichment, and values below 1 indicate depletion.

**Supplementary Figure 4.**
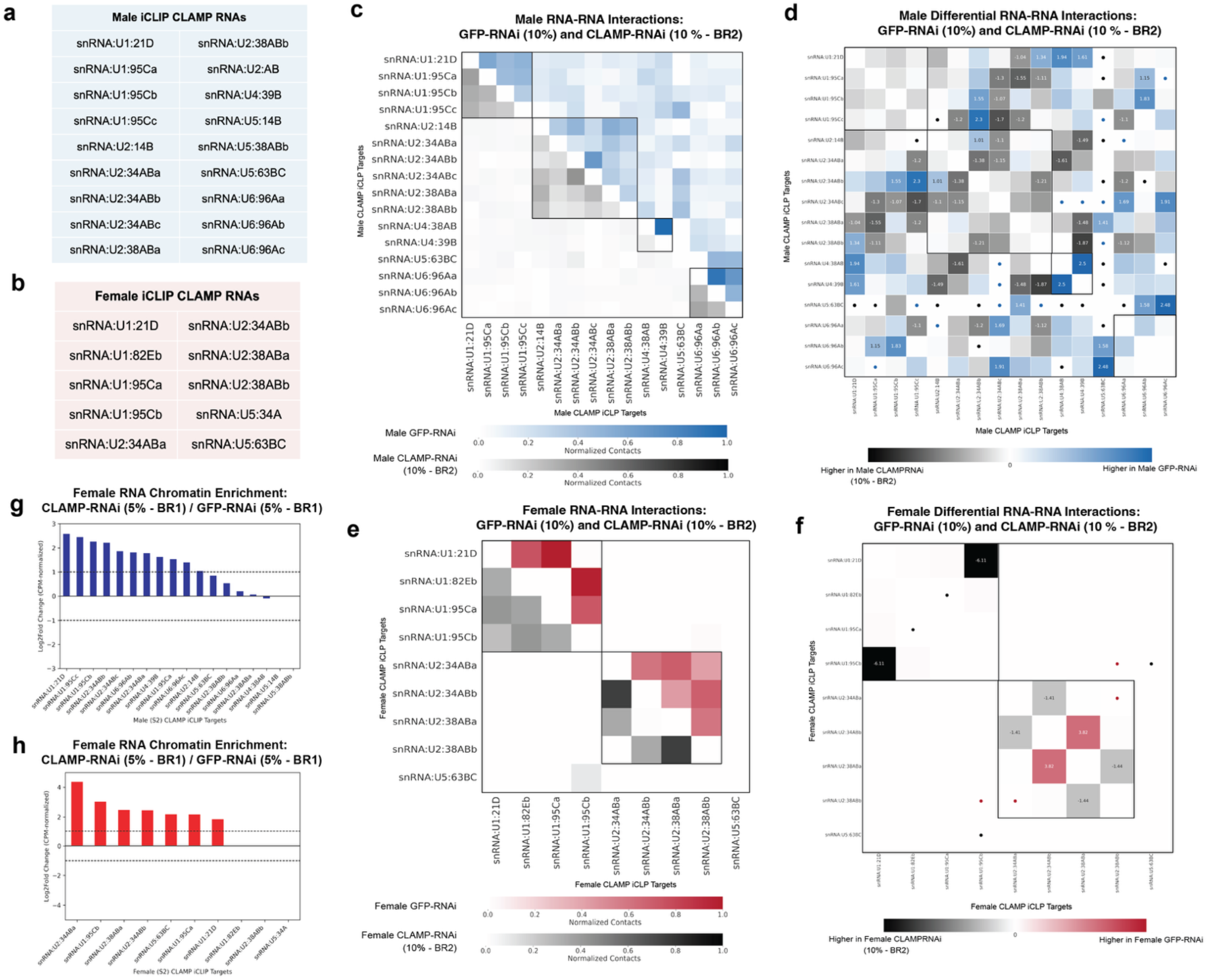
CLAMP depletion is associated with changes in RNA-RNA interactions and chromatin association of CLAMP-bound snRNAs, related to. Figure 3**. (a-b)** snRNAs bound by CLAMP in male (S2, a) and female (Kc167, b) cells identified by iCLIP (Ray et al., 2024). These RNAs were used for all analyses in this figure. **(c)** Normalized RNA-RNA interactions among male CLAMP-bound snRNAs in GFP-RNAi (upper, blue) and CLAMP-RNAi (lower, gray) cells. Boxes outline snRNA classes. **(d)** Differential RNA-RNA interactions between male GFP-RNAi and CLAMP-RNAi cells, shown as normalized log2 fold change. Blue indicates interactions higher in GFP-RNAi, and gray indicates interactions higher in CLAMP-RNAi. Values are shown for interactions with |log2 fold change| > 1. Dots mark interactions detected in only one condition. **(e-f)** Same as (c) and (d), for female cells. In (e), GFP-RNAi is shown in red and CLAMP-RNAi in gray. In (f), red indicates interactions higher in GFP-RNAi, and gray indicates interactions higher in CLAMP-RNAi. **(g-h)** For each snRNA, RNA-DNA contacts were weighted by 2/*n*, where *n* is the number of reads in the RNA’s corresponding cluster. Weighted contacts were normalized to counts per million (CPM) RNA-DNA clusters and summed across the genome. Bars show the log2 fold change of this value in CLAMP-RNAi relative to GFP-RNAi cells in male (g) and female (h) cells. The dotted line represents the 2-fold change threshold. RNA-RNA analyses (c-f) used 10% libraries from biological replicate 2 in Experiment B. Chromatin enrichment analyses (g-h) used 5% libraries from biological replicate 1 in Experiment B.

**Supplementary Figure 5.**
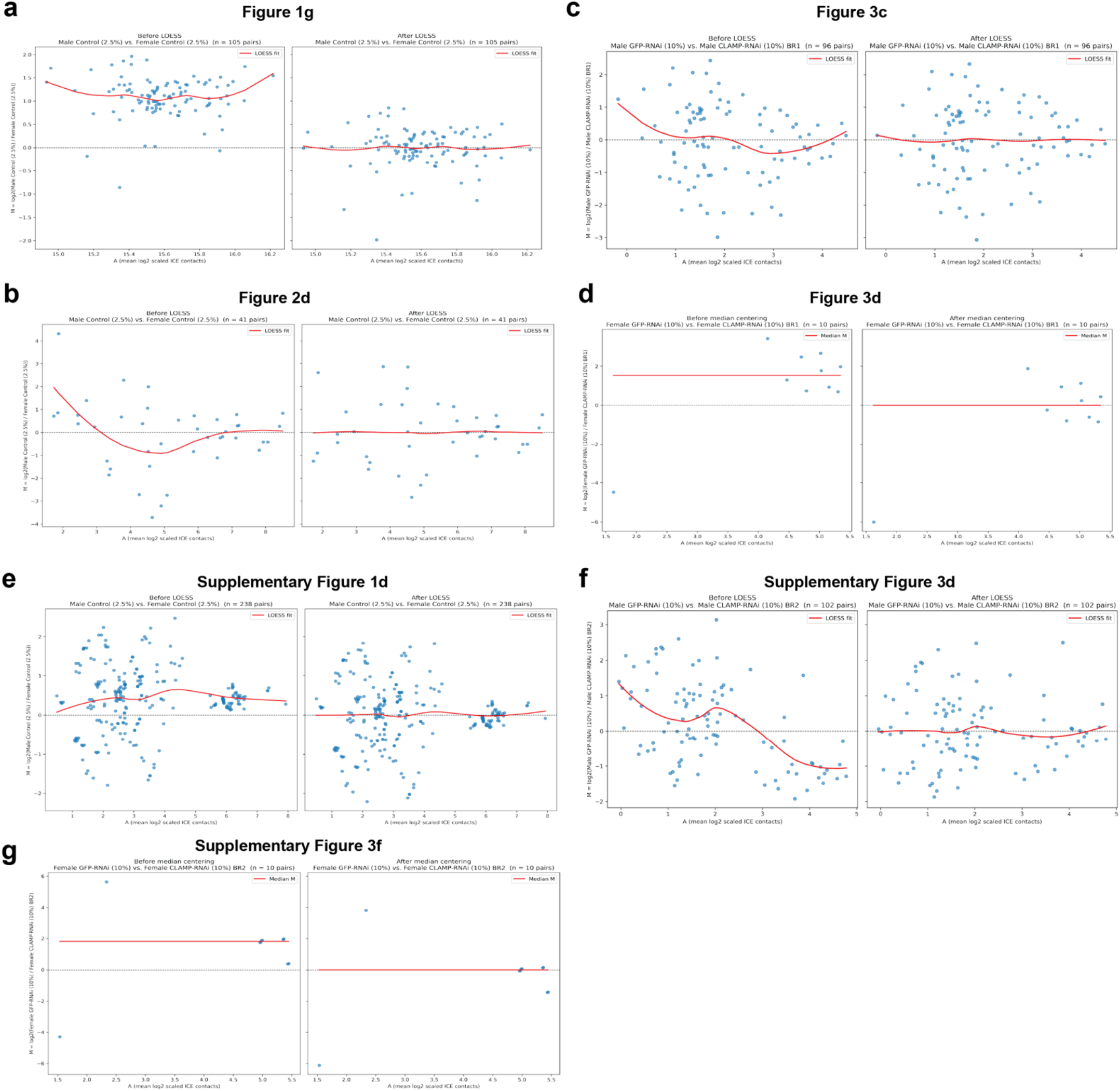
Normalization of differential RNA-RNA interaction analyses. MA plots for each differential analysis, before (left) and after (right) normalization. Each point is an RNA pair. The y-axis shown the log2 fold change in normalized contacts between conditions (M), and the x-axis shown the mean log2 normalized contacts (A). The red line shows the fitted trend. For comparisons with at least 20 RNA pairs, library-specific biases were removed by LOESS normalization (Lun & Smyth 2015; Lun & Smyth 2016). For comparisons with fewer pairs, values were median centered instead, since too few points were available for a reliable LOESS fit. (a) Fig. 1g: Experiment A, male vs. female 2.5% (105 pairs). (b) Fig. 2d: Experiment A, male vs. female 2.5% (41 pairs). (c) Fig. 3c: Experiment B, male GFP-RNAi vs. CLAMP-RNAi, 10%, BR1 (96 pairs). (d) Fig. 3d: Experiment B, female GFP-RNAi vs. CLAMP-RNAi, 10%, BR1, median-centered (10 pairs). (e) Supplementary Fig. 1d: Experiment A, male vs. female 2.5% (238 pairs). (f) Supplementary Fig. 3d: Experiment B, male GFP-RNAi vs. CLAMP-RNAi, 10%, BR2 (102 pairs). (g) Supplementary Fig. 3f: Experiment B. female GFP-RNAi vs. CLAMP-RNAi, 10%, BR2 (10 pairs).

## References

Aguilera, Joseph, Jingyue Duan, Kaitlyn Cortez, et al. 2026. “The CLAMP GA-Binding Transcription Factor Regulates Heat Stress-Induced Transcriptional Repression.” Preprint, March 31. 10.7554/eLife.110960.1.

Alioto, T. S. 2007. “U12DB: A Database of Orthologous U12-Type Spliceosomal Introns.” Nucleic Acids Research 35 (Database): D110–15. 10.1093/nar/gkl796.

Amemiya, Haley M., Anshul Kundaje, and Alan P. Boyle. 2019. “The ENCODE Blacklist: Identification of Problematic Regions of the Genome.” Scientific Reports 9 (1): 9354. 10.1038/s41598-019-45839-z.

Angel, Mor, Eden Fleshler, Mohammad Khaled Atrash, Noa Kinor, Jennifer I. C. Benichou, and Yaron Shav-Tal. 2024. “Nuclear RNA-Related Processes Modulate the Assembly of Cytoplasmic RNA Granules.” Nucleic Acids Research 52 (9): 5356–75. 10.1093/nar/gkae119.

Aquadro, Charles F., Vanessa Bauer DuMont, and Floyd A. Reed. 2001. “Genome-Wide Variation in the Human and Fruitfly: A Comparison.” Current Opinion in Genetics & Development 11 (6): 627–34. 10.1016/S0959-437X(00)00245-8.

Bell, Jason C., David Jukam, Nicole A. Teran, et al. 2018. “Chromatin-Associated RNA Sequencing (ChAR-Seq) Maps Genome-Wide RNA-to-DNA Contacts.” eLife 7 (April): e27024. 10.7554/eLife.27024.

Belmont, Andrew S. 2022. “Nuclear Compartments: An Incomplete Primer to Nuclear Compartments, Bodies, and Genome Organization Relative to Nuclear Architecture.” Cold Spring Harbor Perspectives in Biology 14 (7): a041268. 10.1101/cshperspect.a041268.

Belton, Jon-Matthew, Rachel Patton McCord, Johan Harmen Gibcus, Natalia Naumova, Ye Zhan, and Job Dekker. 2012. “Hi–C: A Comprehensive Technique to Capture the Conformation of Genomes.” Methods 58 (3): 268–76. 10.1016/j.ymeth.2012.05.001.

Bhat, Prashant, Amy Chow, Benjamin Emert, et al. 2024. “Genome Organization around Nuclear Speckles Drives mRNA Splicing Efficiency.” Nature 629 (8014): 1165–73. 10.1038/s41586-024-07429-6.

Blekhman, Ran, John C. Marioni, Paul Zumbo, Matthew Stephens, and Yoav Gilad. 2010. “Sex-Specific and Lineage-Specific Alternative Splicing in Primates.” Genome Research 20 (2): 180–89. 10.1101/gr.099226.109.

Boisvert, François-Michel, Silvana Van Koningsbruggen, Joaquín Navascués, and Angus I. Lamond. 2007. “The Multifunctional Nucleolus.” Nature Reviews Molecular Cell Biology 8 (7): 574–85. 10.1038/nrm2184.

Burge, Christopher B., Richard A. Padgett, and Phillip A. Sharp. 1998. “Evolutionary Fates and Origins of U12-Type Introns.” Molecular Cell 2 (6): 773–85. 10.1016/S1097-2765(00)80292-0.

Camilleri-Robles, Carlos, Raziel Amador, Marcel Tiebe, et al. 2024. “Long Non-Coding RNAs Involved in *Drosophila* Development and Regeneration.” NAR Genomics and Bioinformatics 6 (3): lqae091. 10.1093/nargab/lqae091.

Chen, Li, Dennis J. Lullo, Enbo Ma, Susan E. Celniker, Donald C. Rio, and Jennifer A. Doudna. 2005. “Identification and Analysis of U5 snRNA Variants in *Drosophila*.” RNA 11 (10): 1473–77. 10.1261/rna.2141505.

Drewell, Robert A., Daniel Klonaros, and Jacqueline M. Dresch. 2024. “Transcription Factor Expression Landscape in Drosophila Embryonic Cell Lines.” BMC Genomics 25 (1): 307. 10.1186/s12864-024-10241-1.

Filion, Guillaume J. n.d. Systematic Protein Location Mapping Reveals Five Principal Chromatin Types in Drosophila Cells.

Franke, Axel, and Bruce S. Baker. 1999. “The Rox1 and Rox2 RNAs Are Essential Components of the Compensasome, Which Mediates Dosage Compensation in Drosophila.” Molecular Cell 4 (1): 117–22. 10.1016/S1097-2765(00)80193-8.

Furlong, Eileen E. M., and Michael Levine. 2018. “Developmental Enhancers and Chromosome Topology.” Science 361 (6409): 1341–45. 10.1126/science.aau0320.

Hall, Stephen L., and Richard A. Padgett. 1996. “Requirement of U12 snRNA for in Vivo Splicing of a Minor Class of Eukaryotic Nuclear Pre-mRNA Introns.” Science 271 (5256): 1716–18. 10.1126/science.271.5256.1716.

Hasenson, Sarah E., Ella Alkalay, Mohammad K. Atrash, et al. n.d. The Association of MEG3 lncRNA with Nuclear Speckles in Living Cells.

Hsieh, Tsung-Han S., Assaf Weiner, Bryan Lajoie, Job Dekker, Nir Friedman, and Oliver J. Rando. 2015. “Mapping Nucleosome Resolution Chromosome Folding in Yeast by Micro-C.” Cell 162 (1): 108–19. 10.1016/j.cell.2015.05.048.

Huang, Zheng-hao, Yu-ping Du, Jing-tao Wen, Bing-feng Lu, and Yang Zhao. 2022. “snoRNAs: Functions and Mechanisms in Biological Processes, and Roles in Tumor Pathophysiology.” Cell Death Discovery 8 (1): 259. 10.1038/s41420-022-01056-8.

Huarte, Maite. 2015. “The Emerging Role of lncRNAs in Cancer.” Nature Medicine 21 (11): 1253–61. 10.1038/nm.3981.

Huppertz, Ina, Jan Attig, Andrea D’Ambrogio, et al. 2014. “iCLIP: Protein–RNA Interactions at Nucleotide Resolution.” Methods 65 (3): 274–87. 10.1016/j.ymeth.2013.10.011.

Imakaev, Maxim, Geoffrey Fudenberg, Rachel Patton McCord, et al. 2012. “Iterative Correction of Hi-C Data Reveals Hallmarks of Chromosome Organization.” Nature Methods 9 (10): 999–1003. 10.1038/nmeth.2148.

Jady, B. E. 2003. “Modification of Sm Small Nuclear RNAs Occurs in the Nucleoplasmic Cajal Body Following Import from the Cytoplasm.” The EMBO Journal 22 (8): 1878–88. 10.1093/emboj/cdg187.

Jolly, Caroline, and Subhash C. Lakhotia. 2006. “Human Sat III and Drosophila Hsrω Transcripts: A Common Paradigm for Regulation of Nuclear RNA Processing in Stressed Cells.” Nucleic Acids Research 34 (19): 5508–14. 10.1093/nar/gkl711.

Kalvari, Ioanna, Joanna Argasinska, Natalia Quinones-Olvera, et al. 2018. “Rfam 13.0: Shifting to a Genome-Centric Resource for Non-Coding RNA Families.” Nucleic Acids Research 46 (D1): D335–42. 10.1093/nar/gkx1038.

Kaye, Emily G., Matthew Booker, Jesse V. Kurland, et al. 2018. “Differential Occupancy of Two GA-Binding Proteins Promotes Targeting of the Drosophila Dosage Compensation Complex to the Male X Chromosome.” Cell Reports 22 (12): 3227–39. 10.1016/j.celrep.2018.02.098.

Kharchenko, Peter V., Artyom A. Alekseyenko, Yuri B. Schwartz, et al. 2011. “Comprehensive Analysis of the Chromatin Landscape in Drosophila Melanogaster.” Nature 471 (7339): 480–85. 10.1038/nature09725.

Klonaros, Daniel, Jacqueline M. Dresch, and Robert A. Drewell. 2023. “Transcriptome Profile in Drosophila Kc and S2 Embryonic Cell Lines.” G3 Genes|Genomes|Genetics 13 (5): jkad054. 10.1093/g3journal/jkad054.

Köhler, Alwin, and Ed Hurt. 2007. “Exporting RNA from the Nucleus to the Cytoplasm.” Nature Reviews Molecular Cell Biology 8 (10): 761–73. 10.1038/nrm2255.

Lamond, Angus I., and David L. Spector. 2003. “Nuclear Speckles: A Model for Nuclear Organelles.” Nature Reviews Molecular Cell Biology 4 (8): 605–12. 10.1038/nrm1172.

Lun, Aaron T. L., and Gordon K. Smyth. 2015. “diffHic: A Bioconductor Package to Detect Differential Genomic Interactions in Hi-C Data.” BMC Bioinformatics 16 (1): 258. 10.1186/s12859-015-0683-0.

Lun, Aaron T. L., and Gordon K. Smyth. 2016. “Csaw: A Bioconductor Package for Differential Binding Analysis of ChIP-Seq Data Using Sliding Windows.” Nucleic Acids Research 44 (5): e45–e45. 10.1093/nar/gkv1191.

Mahableshwarkar, Pranav, Jasmine Shum, Mukulika Ray, and Erica Larschan. 2024. “BindCompare: A Novel Integrated Protein–Nucleic Acid Binding Analysis Platform.” Bioinformatics 40 (11): btae668. 10.1093/bioinformatics/btae668.

Malakar, Pushkar, Sudhanshu Shukla, Meghna Mondal, Rajesh Kumar Kar, and Jawed Akhtar Siddiqui. 2024. “The Nexus of Long Noncoding RNAs, Splicing Factors, Alternative Splicing and Their Modulations.” RNA Biology 21 (1): 16–35. 10.1080/15476286.2023.2286099.

Matera, A. Gregory, Rebecca M. Terns, and Michael P. Terns. 2007. “Non-Coding RNAs: Lessons from the Small Nuclear and Small Nucleolar RNAs.” Nature Reviews Molecular Cell Biology 8 (3): 209–20. 10.1038/nrm2124.

Mattick, John S., Paulo P. Amaral, Piero Carninci, et al. 2023. “Long Non-Coding RNAs: Definitions, Functions, Challenges and Recommendations.” Nature Reviews Molecular Cell Biology 24 (6): 430–47. 10.1038/s41580-022-00566-8.

Miao, Hui, Fan Wu, Yu Li, et al. 2022. “MALAT1 Modulates Alternative Splicing by Cooperating with the Splicing Factors PTBP1 and PSF.” Science Advances 8 (51): eabq7289. 10.1126/sciadv.abq7289.

Mir, Mustafa, Michael R. Stadler, Stephan A. Ortiz, et al. 2018. “Dynamic Multifactor Hubs Interact Transiently with Sites of Active Transcription in Drosophila Embryos.” eLife 7 (December): e40497. 10.7554/eLife.40497.

Oksuz, Ozgur, Jonathan E. Henninger, Robert Warneford-Thomson, et al. 2023. “Transcription Factors Interact with RNA to Regulate Genes.” Molecular Cell 83 (14): 2449–2463.e13. 10.1016/j.molcel.2023.06.012.

Ouyang, Jiawei, Yu Zhong, Yijie Zhang, et al. 2022. “Long Non-Coding RNAs Are Involved in Alternative Splicing and Promote Cancer Progression.” British Journal of Cancer 126 (8): 1113–24. 10.1038/s41416-021-01600-w.

Pelletier, Joffrey, George Thomas, and Siniša Volarević. 2018. “Ribosome Biogenesis in Cancer: New Players and Therapeutic Avenues.” Nature Reviews Cancer 18 (1): 51–63. 10.1038/nrc.2017.104.

Pombo, Ana, and Niall Dillon. 2015. “Three-Dimensional Genome Architecture: Players and Mechanisms.” Nature Reviews Molecular Cell Biology 16 (4): 245–57. 10.1038/nrm3965.

Quinn, Jeffrey J., Qiangfeng C. Zhang, Plamen Georgiev, Ibrahim A. Ilik, Asifa Akhtar, and Howard Y. Chang. 2016. “Rapid Evolutionary Turnover Underlies Conserved lncRNA–Genome Interactions.” Genes & Development 30 (2): 191–207. 10.1101/gad.272187.115.

Quinodoz, Sofia A., and Mitchell Guttman. 2022. “Essential Roles for RNA in Shaping Nuclear Organization.” Cold Spring Harbor Perspectives in Biology 14 (5): a039719. 10.1101/cshperspect.a039719.

Quinodoz, Sofia A., Joanna W. Jachowicz, Prashant Bhat, et al. 2021. “RNA Promotes the Formation of Spatial Compartments in the Nucleus.” Cell 184 (23): 5775–5790.e30. 10.1016/j.cell.2021.10.014.

Quinodoz, Sofia A., Noah Ollikainen, Barbara Tabak, et al. 2018. “Higher-Order Inter-Chromosomal Hubs Shape 3D Genome Organization in the Nucleus.” Cell 174 (3): 744–757.e24. 10.1016/j.cell.2018.05.024.

Ray, Mukulika, Ashley Mae Conard, Jennifer Urban, et al. 2023. “Sex-Specific Splicing Occurs Genome-Wide during Early Drosophila Embryogenesis.” eLife 12 (July): e87865. 10.7554/eLife.87865.

Ray, Mukulika, and Julia Zaborowsky. 2026. “A Dual DNA/RNA-Binding Factor Regulates Co-Transcriptional Splicing through Target RNA Interaction and Modulates Splicing Factor Dynamics.” *BiorXiv*, ahead of print, November 1. 10.1101/2024.01.11.575216.

Rivera, Austin, Jou-Hsuan Roxie Lee, Shruti Gupta, et al. 2024. “Traffic Jam Activates the *Flamenco* piRNA Cluster Locus and the Piwi Pathway to Ensure Transposon Silencing and *Drosophila* Fertility.” Preprint, Genetics, August 16. 10.1101/2024.08.15.608167.

Schöfer, Christian, and Klara Weipoltshammer. 2018. “Nucleolus and Chromatin.” Histochemistry and Cell Biology 150 (3): 209–25. 10.1007/s00418-018-1696-3. *Shiny for Python*. n.d. https://shiny.posit.co/py/.

Skene, Peter J., and Steven Henikoff. 2017. “An Efficient Targeted Nuclease Strategy for High-Resolution Mapping of DNA Binding Sites.” eLife 6 (January): e21856. 10.7554/eLife.21856.

Solomon, Mark J., Pamela L. Larsen, and Alexander Varshavsky. 1988. “Mapping proteinDNA Interactions in Vivo with Formaldehyde: Evidence That Histone H4 Is Retained on a Highly Transcribed Gene.” Cell 53 (6): 937–47. 10.1016/S0092-8674(88)90469-2.

Takakuwa, Hiro, Tomohiro Yamazaki, Sylvie Souquere, et al. 2023. “Shell Protein Composition Specified by the lncRNA NEAT1 Domains Dictates the Formation of Paraspeckles as Distinct Membraneless Organelles.” Nature Cell Biology 25 (11): 1664–75. 10.1038/s41556-023-01254-1.

Tarn, Woan-Yuh, and Joan A. Steitz. 1996. “A Novel Spliceosome Containing U11, U12, and U5 snRNPs Excises a Minor Class (AT–AC) Intron In Vitro.” Cell 84 (5): 801–11. 10.1016/S0092-8674(00)81057-0.

Urban, Jennifer A., John M. Urban, Guray Kuzu, and Erica N. Larschan. 2017. “The Drosophila CLAMP Protein Associates with Diverse Proteins on Chromatin.” PLOS ONE 12 (12): e0189772. 10.1371/journal.pone.0189772.

Vitali, Patrice, and Tamás Kiss. 2019. “Cooperative 2′-O-Methylation of the Wobble Cytidine of Human Elongator tRNA^Met^ (CAT) by a Nucleolar and a Cajal Body-Specific Box C/D RNP.” Genes & Development 33 (13–14): 741–46. 10.1101/gad.326363.119.

Wang, Minghui, Yongzhong Zhao, and Bin Zhang. 2015. “Efficient Test and Visualization of Multi-Set Intersections.” Scientific Reports 5 (1): 16923. 10.1038/srep16923.

Wang, Tian, Ezra Wiater, Xinmin Zhang, John B. Thomas, and Marc Montminy. 2021. “Crtc Modulates Fasting Programs Associated with 1-C Metabolism and Inhibition of Insulin Signaling.” Proceedings of the National Academy of Sciences 118 (12): e2024865118. 10.1073/pnas.2024865118.

Yu, Bin, and Ge Shan. 2016. “Functions of Long Noncoding RNAs in the Nucleus.” Nucleus 7 (2): 155–66. 10.1080/19491034.2016.1179408.

Zhang, Huaqun, Vishal Annasaheb Adhav, Audrey C. Kehling, et al. 2026. “Structural Basis for RISC Assembly of Human Argonaute2.” Molecular Cell 86 (11): 2088–2105.e8. 10.1016/j.molcel.2026.04.029.

